# Circulating immune profiling reveals impaired monocyte states and trajectories driving immunosuppression in glioblastoma

**DOI:** 10.64898/2026.01.06.697857

**Authors:** Andrea Scafidi, Tony Kaoma, Claudia Cerella, Eleonora Campus, Kamil Grzyb, Bakhtiyor Nosirov, Frida Lind-Holm Mogensen, Eliane Klein, Raul Da Costa, Alexander Skupin, Frank Hertel, Guy Berchem, Michel Mittelbronn, Antonio Cosma, Beatrice Melin, Simone P Niclou, Petr V Nazarov, Anna Golebiewska, Aurélie Poli, Alessandro Michelucci

## Abstract

Glioblastoma (GBM) is an aggressive and lethal brain tumor marked by profound local and systemic immune dysfunction. Yet, the diagnostic and therapeutic relevance of peripheral impairments remains undefined. To clinically dissect their underlying mechanisms and pathological implications, we combined mass and flow cytometry with single-cell RNA-sequencing of peripheral blood mononuclear cells from GBM patients and healthy donors. GBM blood profiles were characterized by heterogeneous changes in classical monocytes, encompassing expanded, reduced and unchanged subsets, presenting distinct functional states, including antigen-presenting, interferon and metabolic subsets. Additional adaptations included myeloid-derived suppressor cell (MDSC) expansion and loss of non-classical monocytes. Trajectory analyses positioned MDSCs as an intermediate state, in continuum with the metabolic subset. Single-cell RNA-sequencing further showed antigen-presenting monocyte propensity to differentiate into tumor-associated macrophages. Circulating monocytes shared a “GBM-classical monocytic signature” exhibiting low MHC class II expression, altered cell-cell communication and increased anti-inflammatory mediators, such as *IL1R2* and *CD163*. Lastly, lymphocyte alterations included decreased proportions of CD4^+^ T, natural killer (NK) and CD56^+^ T cells, retaining relatively conserved activation profiles, exemplified by up-regulation of alarmins *S100A8/S100A9*. These findings map systemic immune reprogramming in GBM, suggesting new avenues for non-invasive biomarker discovery and therapeutic strategies to restore anti-tumor immunity.

## INTRODUCTION

Glioblastoma (GBM) is the most common and aggressive primary brain tumor accounting for over half of malignant brain tumors in North America and Europe, with an incidence of 2-5 per 100,000 people (*1*). GBM induces profound local and systemic immunosuppression that contributes to disease progression and undermines immune-modulating therapies. Tumor-associated macrophages (TAMs), encompassing microglia (Mg-TAMs) and blood monocyte-derived macrophages (Mo-TAMs), are major local drivers of immune suppression (*2*), displaying high heterogeneity shaped by the tumor and patient-specific characteristics (*3*). Their adaptations within the tumor microenvironment (TME) are reflected in the circulation (*4, 5*), providing a rationale to investigate both compartments for improved patient stratification. Blood-derived immune cells, including myeloid-derived suppressor cells (MDSCs), regulatory T cells and neutrophils contribute to a pro-tumorigenic environment (*6-8*). As in other tumors, systemic immune impairment in GBM features thymic and splenic involution (*9*), T cell retention and dysfunction in the bone marrow (*10*), besides biased hematopoiesis that promotes expansion of myeloid populations, including MDSCs, along with lymphopenia (*11, 12*). In GBM patients, elevated levels of circulating MDSCs, together with reduced natural killer (NK) and CD8⁺ T cells, correlate with poor survival (*13-15*). These findings emphasize the bidirectional interplay between peripheral circulation and the TME (*16*), underscoring the importance of uncovering factors that shape peripheral immune alterations as potential predictors of pathological immune cell infiltration, differentiation and response to immunotherapy.

Mouse GBM models reveal tumor-driven immune perturbations, but single-cell studies in patients remain scarce, focusing mainly on recurrent disease and lacking integrated multi-omic profiling of peripheral immunity. As a result, circulating immune alterations, myeloid trajectories and systemic immunosuppression in primary GBM remain poorly defined. To address this gap, we analyzed peripheral blood mononuclear cells (PBMCs) from a unique cohort of treatment-naive GBM patients using a multimodal single-cell pipeline combining mass cytometry (CyTOF), flow cytometry and single-cell RNA-sequencing (scRNA-seq) to map peripheral immune states and their potential contribution to GBM immunopathology. Circulating immune cells in GBM patients exhibited marked alterations, especially within myeloid cells, CD4^+^ T cells, NK cells and a subset of CD56^+^ T cells displaying mucosal-associated invariant T (MAIT) cell-like features. Myeloid cells exhibited transcriptional reprogramming marked by reduced major histocompatibility complex (MHC) class II-mediated antigen-presentation capacity, increased expression of inflammation-limiting mediators and impaired lineage commitment, revealing disrupted immune trajectories, progression toward tumor-associated macrophages and phenotypic continuity with peripheral MDSCs. Monocytic subsets also differed in proportions, with a substantial loss of non-classical monocytes. Lymphocytes maintained relatively conserved profiles but displayed activation-associated transcriptional signatures. Cell-cell communication analyses inferred immunosuppressive signaling networks, evidenced by major reductions in MHC class II - CD4 interactions.

Overall, our study delineates systemic immune reprogramming in primary GBM and identifies circulating myeloid states with potential to shape the tumor immune landscape, paving the way for the development of novel therapeutic approaches.

## RESULTS

### Peripheral immune cell immuno-profiling uncovers extensive myeloid cell remodeling and reduced lymphoid subsets in GBM

In this study, we investigated whether circulating immune cell profiling in GBM patients can reveal immune trajectories underlying tumor-associated immune heterogeneity, support patient stratification and inform prognostic or therapeutic strategies. To this end, we characterized the peripheral immune cell landscape in GBM, using a multimodal approach (**Fig. 1A**).

**Figure 1.**
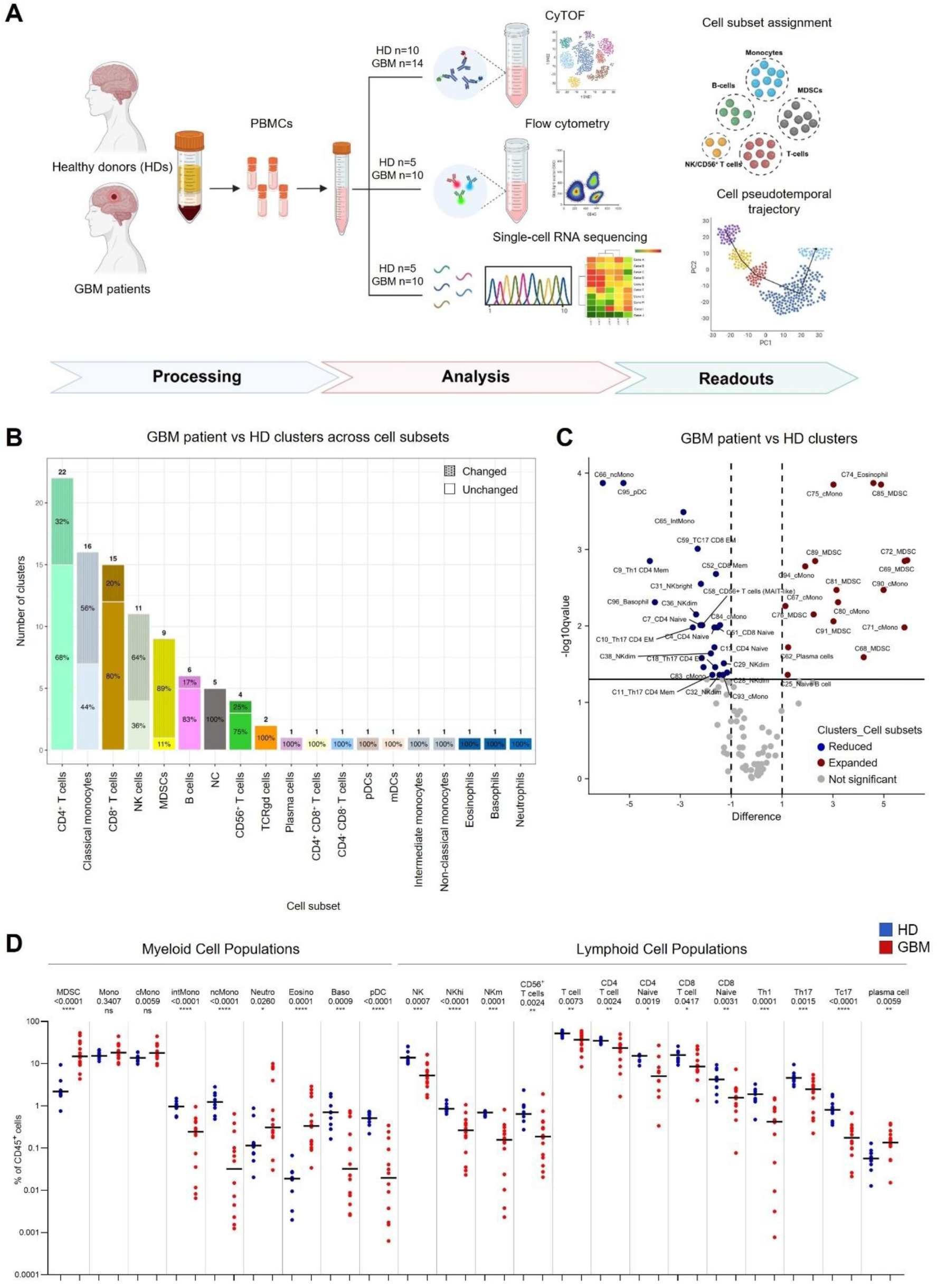
Immune profiling of peripheral blood mononuclear cell alterations in GBM patients. (**A**) Schematic overview of the experimental approach applied in this study (created with BioRender.com). (**B**) Cell subset assignment obtained from unsupervised CyTOF analysis (100-metacluster resolution) using Cell Engine. A threshold-normalized matrix of median cell surface marker expression values (mass intensity units) was used (HDs: n=10; GBM patients: n=14; **Tables S2-S4**). Bar numbers show total clusters belonging to a specific cell type. Percentages represent the fraction of changed (grid pattern) or unchanged cell clusters in GBM patients compared with HDs. (**C**) Volcano plot showing cell subset-assigned clusters identified by Mann-Whitney U test with Benjamini-Hochberg correction, computed in Qlucore. Significantly changed clusters in GBM patients (FDR < 0.05) shown in red when expanded or in blue when reduced (not significant clusters in grey). Vertical dashed lines mark |log_2_FC| > 1. (**D**) Distribution of significantly changed myeloid and lymphoid subsets in HDs (blue) and GBM patients (red) as percentages of CD45^+^ cells. Individual values are shown, and median values are indicated by black lines. The Y-axis is displayed on a logarithmic scale to visualize all subsets together. Statistical analysis: non-parametric two-sided Mann-Whitney U test (p-values: *: < 0.05; **: < 0.01; ***: < 0.001; ****: < 0.0001).

We first performed CyTOF analysis of PBMCs from 14 GBM patients and 10 HDs (**Table S1**, **Fig. S1A**). Unsupervised hierarchical clustering at 100 metaclusters identified all major immune lineages, including myeloid cells, CD4⁺ and CD8⁺ T cells, NK and CD56^+^ T cells, and B cells (**Fig. S1B**). To refine characterization, we classified metaclusters using a threshold-normalized matrix of 33 surface-marker medians, successfully annotating 95 of 100 clusters (**Fig. 1B**, **Tables S2-S3**). Among these, 22 and 15 clusters mapped to CD4⁺ and CD8⁺ T cell subsets, respectively. Seven clusters corresponded to B cell and plasma cell populations, 15 to NK and CD56⁺ T cells and three to CD66b⁺ polymorphonuclear cells. The CD45⁺CD66⁻ myeloid compartment comprised of 28 clusters, including nine with MDSC-like features. Distinct 16 classical monocyte clusters (CD14⁺CD16⁻ cells) emerged. Besides, intermediate monocytes (CD14^+^CD16^+^ cells), non-classical monocytes (CD14^-^CD16^+^ cells) and monocytic dendritic cells (CD14^-^CD16^-^CD11c^+^CD123^-^ cells) each formed a single cluster (**Fig. 1B**, **Table S3**).

We next compared subset abundances across the 100 metaclusters between HDs and GBM patients, revealing altered frequencies in specific immune populations, most prominently diversified in the CD45^+^CD66^-^ myeloid compartment (**Fig. 1B-D**, **Tables S3-S4**, **Fig. S1C-D**). While intermediate and non-classical monocytes were reduced, clusters regrouping classical monocytes exhibited heterogeneous trends of modulation. A subgroup was significantly expanded (clusters 67, 71, 75, 80, 90 and 94), while others were reduced (clusters 83, 84 and 93) or showed no differences (clusters 73, 79, 82, 86, 87, 88 and 92). Although changes in monocytic dendritic cells were not statistically significant, they exhibited a bimodal distribution across GBM patients (**Fig. S1E**), while plasmacytoid dendritic cells were remarkably reduced. Furthermore, eight of the nine MDSC clusters were robustly expanded in GBM patients. Supervised data analysis provided consistent results. Furthermore, multicolor flow cytometry analysis of PBMCs from five HDs and ten GBM patients confirmed a higher frequency of circulating monocytic MDSCs in GBM patients, together with reduced frequencies of intermediate and non-classical monocytes (**Fig. S2-S3**, **Tables S3-S6**).

In addition, the frequency of specific lymphoid clusters was altered, showing an overall reduction in GBM patients, most notably among CD4^+^ and NK cells, as well as in the CD56^+^ T cell subset displaying MAIT-like characteristics. These results were consistent in both unsupervised and supervised CyTOF data analysis (**Fig. 1B-D**, **Fig. S2, Tables S3-S6**). Specifically, we observed a significant decrease in total T cells, accompanied by reduced frequencies of both CD4⁺ and CD8⁺ T cell subsets and their naive sub-compartments from CyTOF analysis. Among CD4⁺ T cells, clusters of Th1 and Th17 subsets were also reduced. While specific CD8⁺ T cell subset frequencies showed limited changes, the Tc17 subset was decreased (**Fig. 1D**, **Table S4**). The T helper Th17/Th1 subsets identified by manual sequential gating were also significantly reduced in GBM patients, including the interferon-γ-producing population (CD4⁺/CD45RA⁻/CD197⁻/CCR4⁻/CCR6⁺/CXCR3⁺/CXCR5⁻). Overall, lymphoid clusters associated with exhausted or regulatory T cells (CD279, TIGIT and/or CD57 expressing CD4^+^ and CD8^+^ T cells) were not modulated or minimally affected (**Fig. S2B**, **Tables S3-S4**). Within the B cell/plasma cell subset, two out of seven clusters were instead significantly increased in GBM patients. Furthermore, NK cells were reduced in both CD56^bright^ and CD56^dim^ subsets as well as CD56^+^ T cells with a MAIT-like phenotype (**Fig. 1D**, **Fig. S2-S3**, **Tables S3-S6**).

Altogether, these data reveal specific alterations in the peripheral immune cell composition of GBM patients, indicating a pronounced and more heterogeneous modulation of the myeloid compartment, in addition to an overall decrease in CD4^+^ T and NK/CD56^+^ T cell subsets.

### Frequency-based macrocluster profiling of circulating myeloid cells reveals phenotypic shifts and disrupted trajectories in GBM

The myeloid lineage shows marked developmental plasticity, generating monocytic subsets with distinct functions. This balance is disrupted in GBM, favoring expansion of immunosuppressive myeloid cells, including MDSCs (*17*). In parallel, we showed that intermediate and non-classical monocytes are uniformly reduced, whereas classical monocyte clusters display heterogeneous changes, with subsets expanding, contracting or remaining stable. To assess whether this heterogeneity reflects altered myeloid differentiation or activation trajectories, phenotypic dynamics across subsets were evaluated using trajectory analysis. To this aim, we consolidated the classical monocyte-like clusters identified by unsupervised CyTOF analyses into broader macroclusters (MCs), based on their modulation pattern in GBM: MC-reduced, MC-unchanged and MC-expanded (**Fig. S4A**, **Table S3**). MDSCs and intermediate/non-classical monocytes were included as distinct MCs to assess their potential phenotypic continuity with the other MCs (**Fig. S4A-B**). Phenotypic profiles of all MCs were then examined using surface markers with detectable expression in the threshold-normalized matrix **(Table S2)**, including regulators of lineage differentiation and activation markers. Expression of HLA-DR, CD45, CD33, CD14 and CD11c progressively decreased from MC-reduced to MC-expanded (**Fig. 2A-B** and **Fig. S4C**), suggesting a gradual loss of antigen-presenting capacity and incomplete myeloid maturation in GBM. Notably, MC-expanded displayed phenotypes closely resembling MDSCs, despite not fully matching the canonical CD33^+/^HLA-DR^dim^ profile. These MC-expanded also showed reduced CD123 (interleukin 3 receptor, IL-3R) expression, indicating impaired IL3 responsiveness, a pathway relevant for impaired anti-tumor activity (**Fig. 2A** and **Fig. S4C**). Multicolor flow cytometry confirmed an overall decrease in surface marker expression in CD14^+^CD16^-^ myeloid cells from GBM patients with similar decrease of HLA-DRA, CD45, CD33 and CD11c observed across all the monocytic subsets (CD14^+^CD16^-^, CD14^+^CD16^+^ and CD14^-^CD16^+^ cells) (**Fig. S5**).

**Figure 2.**
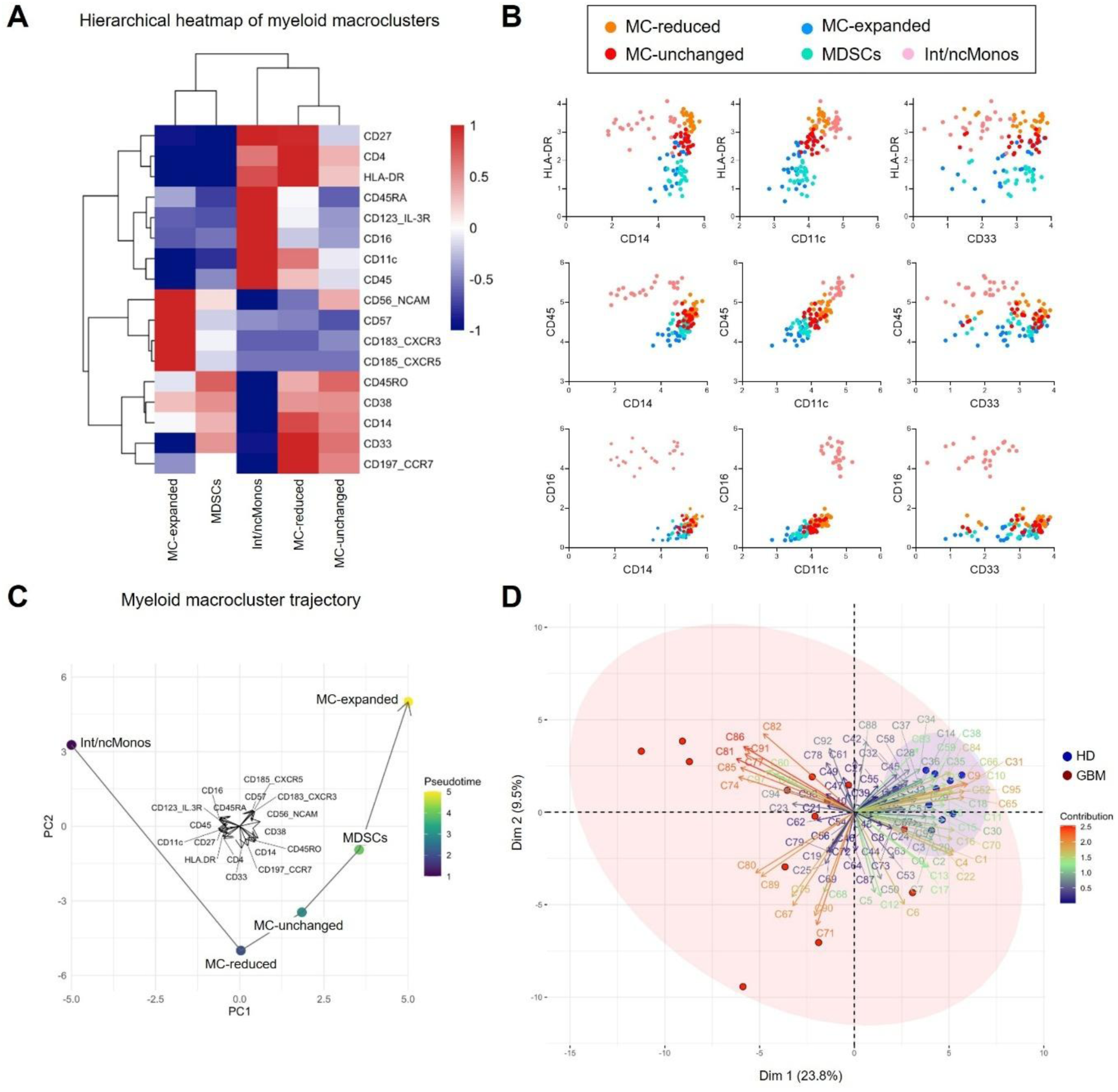
Circulating myeloid immune cells display altered phenotypes and trajectories in GBM patients. (**A**) Hierarchical heatmap of the five defined macroclusters, showing distinct signatures based on myeloid differentiation markers and markers expressed in the threshold-normalized based matrix (color bar denotes Z-score value scale of median expression values). Kruskal-Wallis test shows significant differences in all markers (p-value < 0.0001) except CD57, CXCR3, and CXCR5. (**B**) Median expression trends of selected markers across the five macroclusters. MC: macrocluster; Int/ncMonos: intermediated/non-classical monocytes. (**C**) Trajectory analysis of the five macroclusters projected in the principal component (PC) space. Color bar denotes pseudotime. (**D**) Biplot combining PCA coordinates with unsupervised-defined cluster contributions showing PCA scores of the explanatory variables as vectors (arrows) and clusters of each group (red: GBM patients, blue: HDs) of the first (X-axis) and second (Y-axis) PCs. Arrow direction indicates the orientation of each cluster in the PC space, while arrow length and color represent the magnitude of each cluster contribution to the PCs. Points are individual samples and those located closer to a cluster arrow have higher representation of that cluster features. Colored concentration ellipses (size determined by a 0.95-probability level) show the observations assembled by groups (blue: HDs, red: GBM patients).

Trajectory analyses revealed a continuum linking intermediate/non-classical monocytes to MC-reduced and MC-unchanged, with the latter positioned proximal to MDSCs (**Fig. 2C**). MC-expanded aligned downstream of this trajectory and emerged from MDSCs, displaying combined modulation of HLA-DR, CD45, CD33, CD14 and CD11c, alongside increases in CXCR3/5, CD57 and CD56. These patterns imply that MDSCs represent transitional stages that precede the development of MC-expanded classical monocyte subsets (**Fig. 2C**). Principal component analysis (PCA) confirmed phenotypic proximity of MC-expanded subsets and MDSCs. Moreover, HDs showed close clustering, whereas GBM patients appeared widely distributed across the PCA space, reflecting inter-patient variability mainly associated with MC-expanded populations and MDSCs (**Fig. 2D**).

Overall, these findings indicate that GBM induces extensive myeloid reprogramming, with expansion of immunosuppressive monocytic populations and a differentiation continuum in which MDSCs occupy intermediate transitional states.

### GBM patients show prominent transcriptional changes in monocytes and relatively conserved lymphocytic states compared with healthy donors

To link the phenotypic alterations identified by CyTOF and flow cytometry to underlying transcriptional programs, we next conducted scRNA-seq analysis of PBMCs from ten GBM patients and five HDs (**Fig. 1A**, **Table S1**). UMAP embedding of samples from GBM patients and HDs documented a distribution of peripheral immune cells across 12 non-overlapping clusters (**Fig. 3A**). Using the Atlas Azimuth as reference to guide cell type annotation (*18*) and verifying the levels of expression of specific cell type markers (**Fig. 3B**), we identified major immune cell types defined as T cells (cluster 1), monocytes and dendritic-like cells (clusters 2-7), NK cells (cluster 8), B cells (cluster 9 and 10), platelets (cluster 11) and precursor cells (cluster 12) (**Fig. 3A-B**). Following further refinement of T cells into CD4^+^ T cells, CD8^+^ T cells and other T cells, we quantitatively verified the cluster composition by cell types clearly showing the identities of the main groups (**Fig. 3C**). Next, we compared the composition of the clusters between GBM patients and HDs and observed that while lymphocytic clusters were composed of cells from HD and GBM blood (clusters 1 and 8-10), monocytic clusters mainly contained cells from either GBM patients (clusters 2, 3 and 5-7) or HDs (cluster 4) (**Fig. 3D**), suggesting more prominent transcriptional differences within the monocytic compartment compared to lymphocytic cells between GBM patients and HDs. We further observed that while monocytes from HDs showed similar transcriptomic profiles (cluster 4), monocytes from GBM patients were distributed across shared (clusters 2, 3) or patient-specific clusters (clusters 5-7) (**Fig. 3E**), indicating a certain degree of variability in their transcriptional changes among patients.

**Figure 3.**
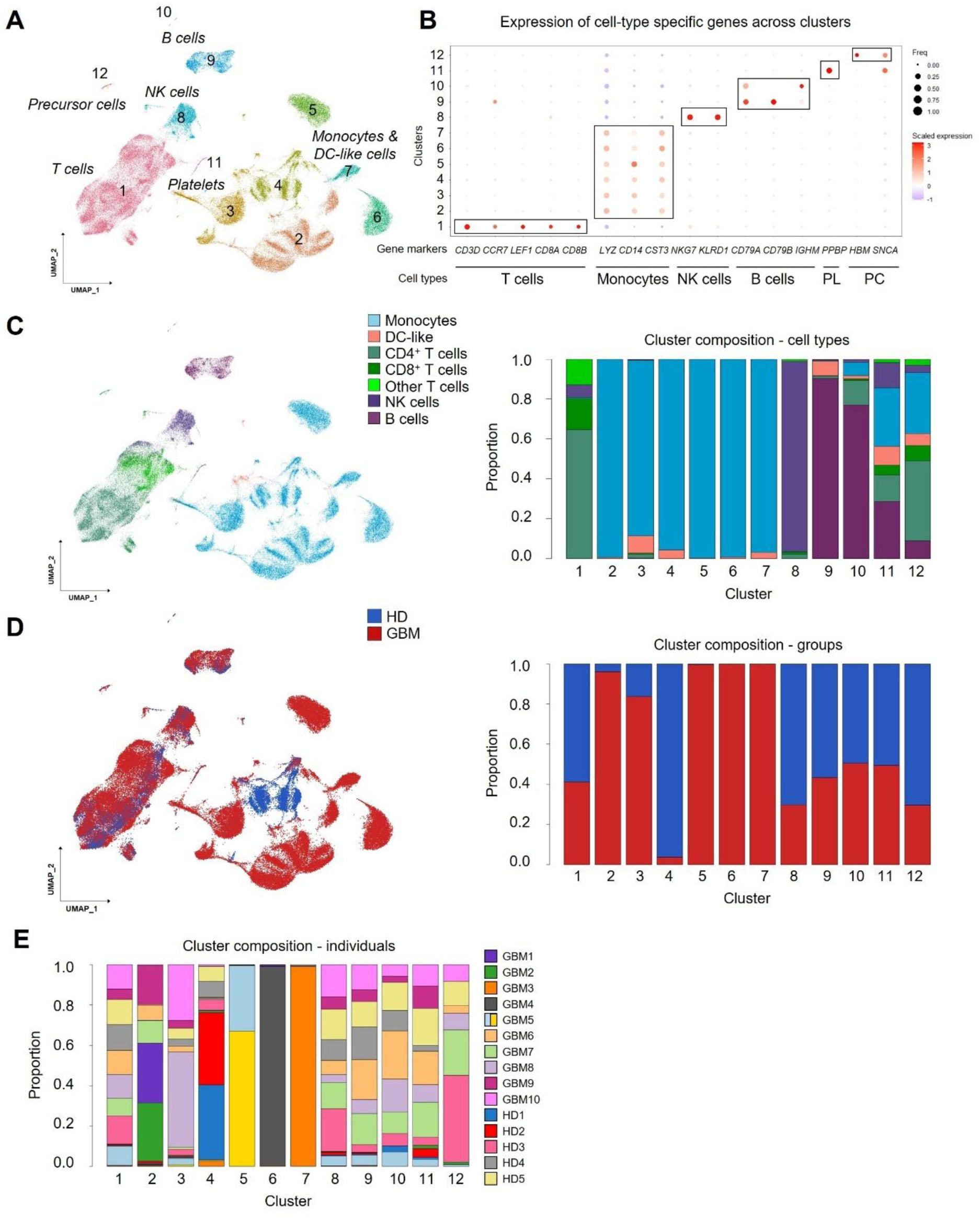
GBM patients show prominent monocytic cell transcriptional changes and relatively conserved lymphocytic cell profiles compared with healthy donors. (**A**) UMAP showing PBMCs from GBM patients and HDs segregating into 12 different clusters. (**B**) Dot plot depicting the expression of cell-type specific genes across different clusters (adjusted p-value < 0.05, Avg_log_2_FC ≥ 0.25; **Table S7**). (**C**) Left: UMAP showing cell type assignation from Azimuth: monocytes (blue), dendritic cell (DC)-like (pink), CD4^+^ T cells (dark green), CD8^+^ T cells (bright dark green), other T cells (light green), NK cells (purple) and B cells (magenta). Right: Bar chart showing the composition of cell types in each cluster. (**D**) Left: UMAP depicting PBMCs from GBM patients (in red) and HDs (in blue). Right: Bar chart showing the composition of GBM patient-and HD-derived cells across the 12 clusters. (**E**) Bar chart showing the individual contribution to the PBMC clusters.

These data show that scRNA-seq of PBMCs from GBM patients and HDs resolves the major blood immune cell populations and reveals that GBM-associated transcriptional changes are more pronounced in monocytes than in lymphoid cells.

### Monocytes exhibit diverse transcriptional programs, including antigen-presenting cell, MDSC-like, metabolic, non-classical, interferon and ribosomal signatures, with varying proportions between GBM patients and healthy donors

Following the observation that monocytes exhibit prominent transcriptional alterations in GBM patients, we sought to further resolve the monocytic clusters (**Fig. 3A**). After normalization of monocytes (*19*), we first discriminated classical from non-classical monocytes (**Fig. 4A**). Within the classical monocytic cluster, we identified five discrete subsets. These subsets were subsequently characterized based on the genes significantly upregulated in each subset relative to the others (log_2_FC ≥ 0.25, adjusted p-value < 0.05) (**Fig. 4B**, **Table S7**). PCA of these distinguished monocytes revealed that transcriptional weighted average proportion variance was primarily driven by cell subtypes (weighted average: 0.397) with minimal contribution from sample and batch, as well as group and dexamethasone treatment (**Fig. S6A**). We detected similar patterns when separately analyzing HDs and GBM patients (**Fig. S6B-C**). Cluster 1, designated as antigen-presenting cell (APC) monocytic group, showed the highest enrichment in genes encoding MHC class-II components (e.g. *HLA-DPB1, HLA-DRA* and *CD74*), while cluster 2 was enriched with MDSC-marker genes (e.g. *S100A8, S100A12* and *S100A9*), lacking antigen presentation markers. We verified the identity of the recognized MDSC-like cluster using literature-based MDSCs and HLA-DR^low^S100A^high^ transcriptional signatures (*20-22*) (**Fig. S6D**). Cluster 3, named as the metabolic monocytic subset, was characterized by genes related to metabolic processes, such as oxidative phosphorylation (e.g. *MT-CYB, PDK4* and *BACH1*), while cluster 4 was represented by the expression of the prototypical non-classical monocytic marker *FCGR3A* (encoding CD16). Lastly, cluster 5 was enriched with genes related to interferon responses and immune cell activation against viruses (e.g. *MX1, IFI44L* and *ISG15*), while cluster 6 showed the induction of genes related to ribosomal activity and cytoplasmic translation (e.g. *RPS26, RPS18* and *RPS27*) (**Fig. 4B-C**, **Fig. S6E, Table S8**). To further determine the transcription factors (TFs) whose activation specifies each individual cluster, we applied single-cell regulatory network inference and clustering (SCENIC) (*23*).

**Figure 4.**
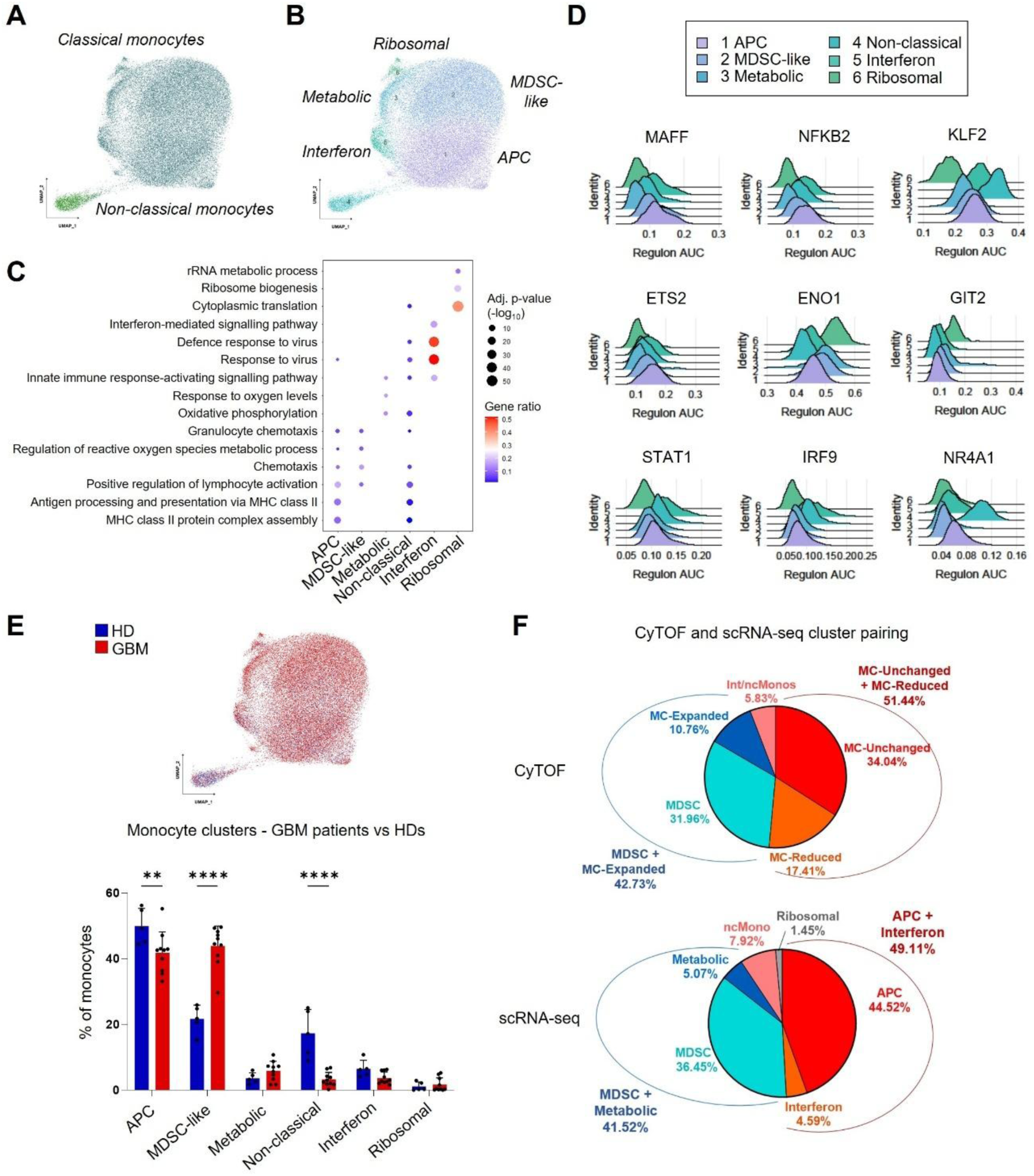
GBM patients and healthy donors display discrete transcriptional monocytic subsets with differing proportions. (**A**) UMAP showing re-clustered monocytes from UMAP depicted in Fig. 3C discriminating classical and non-classical monocytes. (**B**) UMAP showing five distinct classical monocytic clusters: APC, MDSC-like, metabolic, interferon and ribosomal (**Table S7**). (**C**) Dot plot depicting main biological processes defining the identified monocytic clusters (**Table S8**). Circle diameter depicts -log_10_ (adjusted p-value) and the color denotes gene ratio. (**D**) Ridge plots showing selected regulons across monocytic subsets (**Table S9**). (**E**) Top: UMAP showing monocytic cells from HDs (blue) and GBM patients (red). Bottom: bar chart displaying the percentage of GBM patient- and HD-derived cells in each monocytic cluster. Unpaired t-test (n=10 GBM patients, n=5 HDs; p-values: **: < 0.01; **** < 0.0001). (**F**) Correspondence between CyTOF and scRNA-seq clusters: MDSCs and intermediate/non-classical monocytes in the CyTOF analysis matching scRNA-seq MDSC and non-classical clusters; combined MDSC/expanded monocytic clusters in CyTOF correlating with MDSC/metabolic clusters from scRNA-seq analysis; combined unchanged/reduced monocytic clusters in CyTOF overlapping with APC/interferon clusters from scRNA-seq analysis. MDSCs: light blue; intermediate/non-classical monocytes: pink; expanded/metabolic monocytic clusters: dark blue; unchanged monocytic/APC clusters: red; reduced monocytic/interferon clusters: orange; ribosomal cluster: grey.

Here we identified APC, non-classical and interferon monocytes characterized by the activation of TFs related to inflammatory responses (e.g. MAFF, NFKB2, KLF2 and ETS2), while MDSC-like, metabolic and ribosomal clusters were mainly distinguished by the induction of TFs negatively regulating immune activation (e.g. ENO1 and GIT2). Evidently, non-classical and interferon monocytes were associated with interferon responses (e.g. STAT1 and IRF9) (**Fig. 4D**) (**Table S9**). We further verified the relevance of the defined monocytic signatures within the transcriptional programs described in GBM PBMCs by Miller and colleagues (*24*). The APC signature was highly represented in the “*systemic inflammatory*” program, whereas the MDSC-like signature was enriched in the “*CD163^+^ monocyte*” program. As anticipated, non-classical markers were most prominent in the “*CD16^+^ monocyte*” program, while the interferon signature was associated with the “*interferon response*” program. In contrast, the metabolic and ribosomal signatures did not clearly align with the 11 predefined programs, suggesting a peculiar composition (**Fig. S6F**). To exclude potential contamination by other cell types within our defined subsets, we confirmed that all clusters were effectively enriched for the “*CD14^+^ monocyte*” program, with lower expression in the non-classical group (**Fig. S6F**).

All the clusters were represented across the two groups, though GBM patients showed a significant reduction of APC and non-classical monocytes together with an increase of MDSC-like cells compared with HDs (**Fig. 4E**), in line with CyTOF (**Fig. 1C-D**) and flow cytometry (**Fig. S3B**) analyses. Lastly, we assessed correspondence between the myeloid clusters identified by CyTOF and scRNA-seq. Frequencies of MDSCs and intermediate/non-classical monocytes were highly concordant, with the combined MDSC and metabolic scRNA-seq clusters matching the MDSC and MC-expanded groups defined by CyTOF. Conversely, although the APC and interferon clusters differed slightly in their individual proportions, their combined frequencies corresponded closely to the MC-unchanged and MC-reduced groups, indicating subtle transitional states (**Fig. 4F**).

In line with cytometry analyses, these results indicate that GBM patients exhibit a reduction in functional monocytic clusters, including APC and non-classical monocytes, accompanied by an increase in immunosuppressive MDSC-like cells.

### Monocyte subsets follow distinct developmental trajectories in the circulation and during differentiation into tumor-associated macrophages

To characterize the relationships among the identified monocytic clusters, we conducted a pseudotime analysis reconstructing their developmental trajectories using TSCAN algorithm (*25*). Similarly to the trajectories identified by CyTOF, we uncovered distinct apparent developmental paths originating from APC monocytes: the first progressing to ribosomal through MDSC-like and metabolic monocytic clusters, the second evolving towards the interferon subset and the third differentiating into non-classical monocytes (**Fig. 5A**). The first developmental path was enriched in GBM patients according to the corresponding increase of the MDSC-like cluster compared to HDs and was characterized by the upregulation of prototypical MDSC markers, such as *S100A8, S100A9* and *S100A6*, together with the decrease of immune activation markers, including *NFKBIA, IL1B* and *CD74*. The interferon-associated pathway was characterized by the induction of stereotypical markers related to myeloid cell stimulation via interferons, including *IFI44, IFITM3* and *STAT1*, along with the downregulation of inflammatory and APC markers (e.g. *IL1β, CD74* and *HLA-DRA*). The non-classical monocyte pathway exhibited enhanced expression of its defining markers (e.g. *CDKN1C, LST1* and *MS4A7*) and downregulation of classical monocytic-specific genes (e.g. *VCAN, LYZ* and *CD14*) (**Fig. 5B, Table S10**).

**Figure 5.**
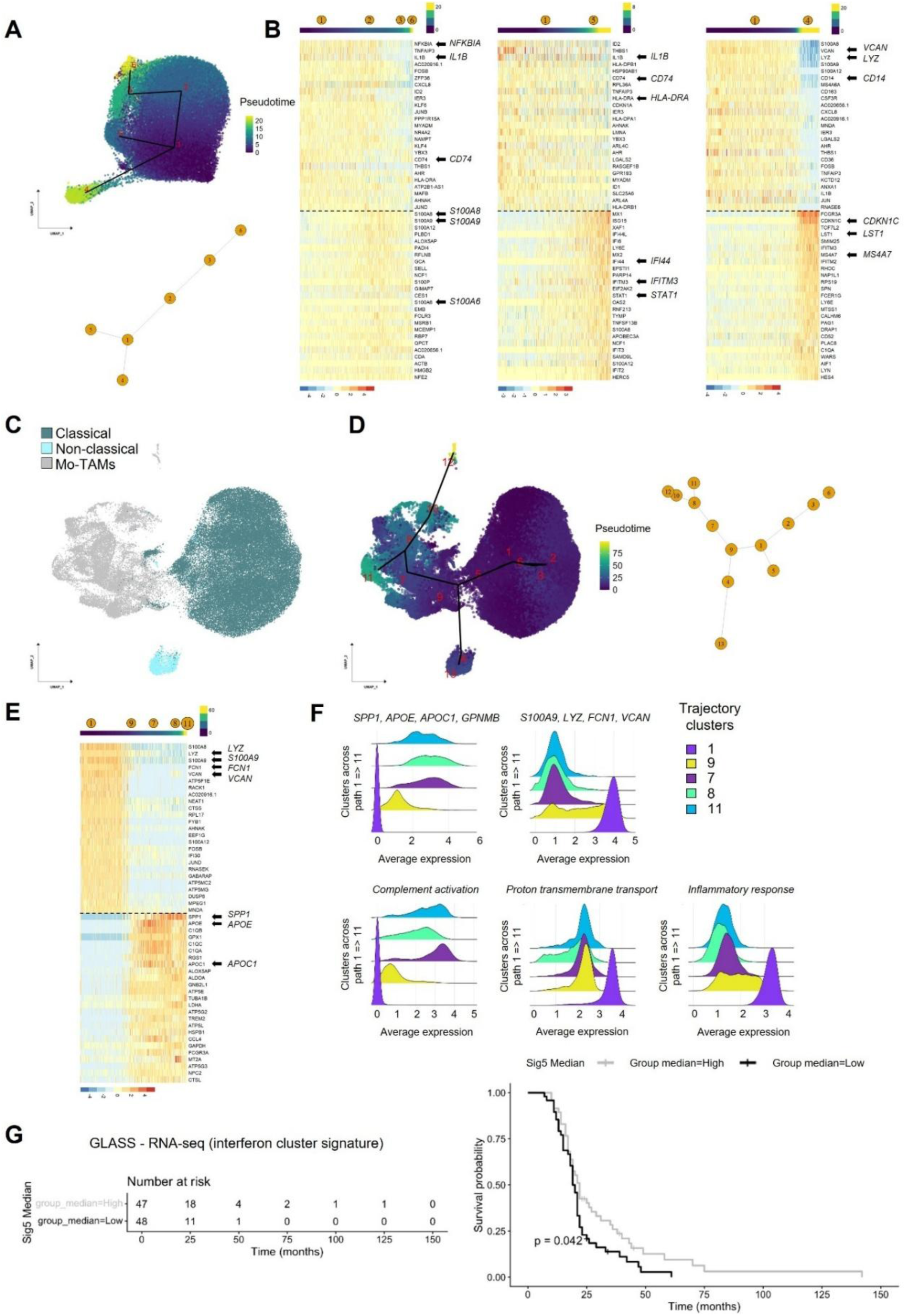
Monocyte subsets follow discernible developmental trajectories in circulation and during differentiation into tumor-associated macrophages. (**A**) Top: UMAP depicting trajectories across monocytic subsets. Color bar represents pseudotime. Bottom: schematic showing the trajectories identified via TSCAN algorithm (cluster 1 as origin). (**B**) Heatmap showing the top 20 downregulated (top) and upregulated (bottom) genes along the trajectory from cluster 1 to 6 (left), from cluster 1 to 5 (center) and from cluster 1 to 4 (**Table S10**). Upper color bar represents pseudotime, lower color bar denotes scaled gene expression level. (**C**) UMAP showing blood classical monocytes (dark blue) and non-classical monocytes (light blue) from HDs and GBM patients integrated with intratumoral Mo-TAMs (grey) identified in Yabo et al. (*3*). (**D**) Left: UMAP showing trajectories across blood monocytic subsets and Mo-TAMs. Color bar represents pseudotime. Right: schematic illustrating pseudo-times calculated with TSCAN algorithm (cluster 1 as origin). (**E**) Heatmap showing the top 20 downregulated (top) and downregulated (bottom) genes along the trajectory from cluster 1 to cluster 11 (**Table S11**). Upper color bar represents pseudotime, lower color bar denotes scaled gene expression level. (**F**) Top: ridge plots showing the average expression level of prototypical TAM markers (*SPP1, APOE, APOC1* and *GPNMB*) and monocytic-specific genes (*S100A9, LYZ, FCN1* and *VCAN*) along the trajectory from cluster 1 to 11. Bottom: ridge plots representing the average expression level of GO specific gene signatures along the trajectory from cluster 1 to 11: complement activation (GO:0006958; *C1QB, C3, C1QA, C1QC*), proton transmembrane transport (GO:1902600; *MT-ATP6, RNASEK, MT-CO1, MT-CO2, MT-CO3, MT-ND3, ATP6V0C, ATP5F1E, MT-ND1*) and inflammatory response (GO:0006954; *ANXA1, NAMPT, CYBB, TXNIP, S100A12, FOS, LYZ, S100A9, S100A8*). (**G**) Kaplan-Meyer curve of overall survival for primary IDH-WT GBMs in the GLASS cohort (*27*) stratified according to the interferon monocytic signature (cluster 5). A two-sided log-rank test was used to calculate the p-value.

Next, given that TAMs can arise from circulating monocytes, we aimed to determine whether any of these monocytic subsets display a higher propensity to differentiate into Mo-TAMs within the TME. To address this, we integrated our GBM blood monocyte dataset with published scRNA-seq data from GBM tumors, selecting Mo-TAMs detected in patient samples for trajectory analysis (**Fig. 5C, Table S11**) (*3*). We identified two Mo-TAM differentiation paths (pseudotime ending with cluster 11 and 12) exclusively arising from APC classical monocytes (cluster 1), with no contribution from other subsets (**Fig. 5D**), reflecting a greater propensity of this subset to be recruited into the tumor tissue and transition towards Mo-TAMs. As expected, both trajectories showed progressive upregulation of prototypical TAM markers (e.g. *SPP1, APOE, APOC1* and *GPNMB*), together with downregulation of monocytic genes (e.g. *S100A9, LYZ, FCN1* and *VCAN*). Alongside with the gradual up-regulation of genes associated with “complement activation” (e.g. *C1QB, C1QC, C1QA* and *C3*), genes related to “proton transmembrane transport” (e.g. *ATP5F1E* and *RNASEK*) and “inflammatory response” (e.g. *TXNIP* and *FOS*) were down-regulated in both trajectories towards Mo-TAMs (**Fig. 5E-F, Fig. S7A-C, Table S11**). Next, we further took advantage of the described transcriptional identity and activity programs in the TME (*24*) to assess their relevance within our defined monocytic subsets. The non-classical monocytic cluster scored low for the “*monocyte*” program. Additionally, APC and interferon clusters showed high scores for the “*systemic inflammatory*” program, whereas non-classical and interferon subsets scored high for the “*interferon response*” signature (**Fig. S7D**), highlighting the relevance of TME signatures to define peripheral monocytic populations.

In this context, we examined whether our monocytic gene signatures were linked to overall survival. To this end, we used the top up-regulated genes from each monocytic-associated cluster (log2FC ≥ 0.25, adjusted p-value < 0.05) to score IDH-WT GBMs within the Glioma Longitudinal Analysis Consortium (GLASS) dataset. We found that the interferon-related monocytic program was significantly associated with improved overall survival, whereas no such association was observed for the other monocytic signatures (**Fig. 5G**).

Lastly, we spatially determined the expression levels of classical APC monocyte genes and Mo-TAM genes that were modulated across the monocytic differentiation into Mo-TAMs (**Fig. 5E, Fig. S7A, Fig. S7C**) and exclusively expressed by non-tumoral cells taking advantage of the Ivy Glioblastoma Atlas project (*26*). APC monocytic genes were predominantly expressed in the perivascular areas and their expression decreased from the tumor core towards the margins, thus suggesting that perivascular areas correspond to zones of extravasation of blood monocytes into the tissue. On the other hand, the Mo-TAM-specific gene signature showed higher expression across the pseudopalisading and perinecrotic regions, while the marginal areas were dominated by Mg-TAM specific transcripts, such as *P2RY12* and *TMEM119* (**Fig. S7E**).

Taken together, APC monocytes generate discrete circulating subsets that preferentially infiltrate tumors and differentiate into defined Mo-TAMs, yielding both immunosuppressive monocytes and tumor-supportive macrophages. Notably, the interferon program determines overall survival.

### Classical monocytes in GBM patients show diminished MHC class II presentation capacity together with upregulation of inflammation-limiting mediators

Next, we analyzed differentially expressed genes (DEGs) between GBM patients and HDs across the identified classical monocytic clusters to determine the effect of GBM on the specific subsets. All the clusters, except for the ribosomal group, possibly because of the low number of cells, exhibited a consistent number of DEGs (APC: 280; MDSC-like: 243; metabolic: 139; interferon: 87; (|log_2_FC| ≥ 0.25, adjusted p-value < 0.05) (**Fig. S8A, Table S12**). When compared with HDs, these identified four clusters in GBM patients showed a common transcriptional signature that we named “glioblastoma-classical monocytic signature” (GBM-CMS), characterized by the upregulation of 29 genes mainly associated with monocytic anti-inflammatory functions, such as the top enhanced genes *IL1R2*, *CD163* and *FKBP5*, and the concomitant downregulation of 6 genes mostly related to MHC class II protein complex, including *HLA-DRB5* and *HLA-DQB1*. Genes exhibiting distinct modulation within these clusters were linked, for instance, to extracellular matrix organization and cell adhesion, including *VCAN, LGALS1* and *ICAM1*, most prominently in the APC cluster, whereas genes such as *MAFB, ETS2*, *IRF1* and *KLF2*, associated with myeloid differentiation and transcriptional regulation, were primarily down-regulated in the MDSC-like group (**Fig. 6A, Fig. S8B, Table S13**). Flow cytometry confirmed increased proportion of CD163^+^ cells in GBM-derived CD14^+^CD16^-^ classical monocytes (**Fig. 6B**) along with decreased expression levels of CD11c (encoded by *ITGAX*), HLA-DR and CD4 proteins (**Fig. S5**), thus confirming their enhanced immunosuppressive traits and reduced APC capacities. According to the common GBM signature, GO terms were similar across the various classical monocytic subsets (**Table S13**), with terms related to downregulated genes mainly associated to “regulation of T cell activation” in the APC and MDSC-like clusters, while terms related to “MHC protein complex assembly” mostly linked to the metabolic and interferon groups (**Fig. S8C**). Further, we identified seven TFs commonly downregulated across the four monocytic clusters, including the retinoic acid receptor RARA and RUNX3, key regulators of monocyte differentiation toward distinct immunosuppressive programs (*28, 29*). Specific upregulated regulons in the APC cluster, such as PICK1, induced in macrophages under pro-inflammatory conditions restraining their prototypical activation (*30*), as well as inactivation of NFKB2 (p52), an important subunit of the non-canonical NF-κB signaling (*31, 32*), in the APC and metabolic clusters, further show their impaired immune responses (**Fig. 6C**).

**Figure 6.**
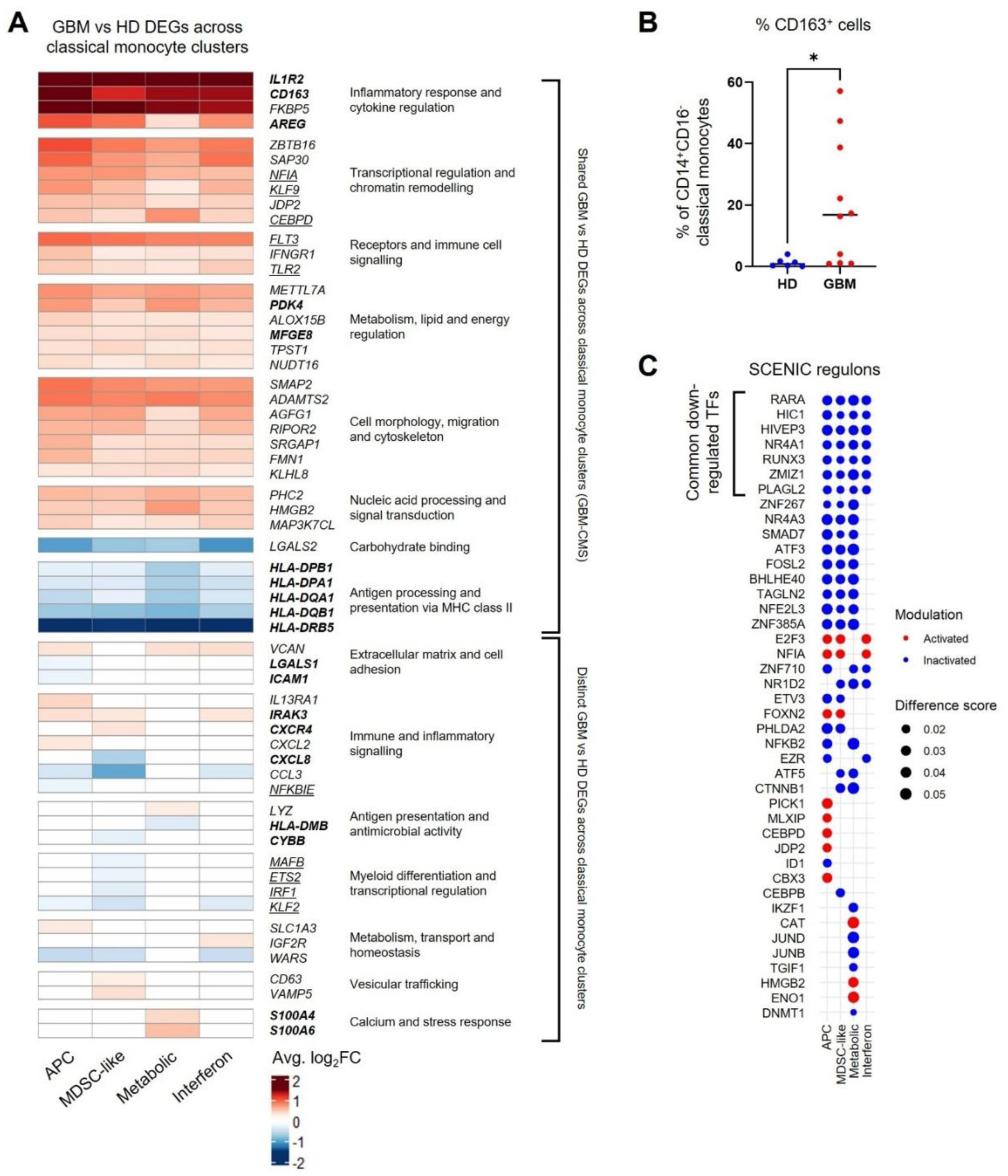
Classical monocytes in GBM patients show reduced ability to present antigens via MHC class II molecules alongside upregulation of genes associated with immune evasion pathways. (**A**) Heatmap relative to DEGs comparing GBM patients (n=10) and HDs (n=5) (adjusted p-value < 0.05, |Avg_log2FC| ≥ 0.25; **Table S12**) showing shared (GBM-CMS) and distinct genes across APC, MDSC-like, metabolic and interferon monocytic clusters according to the Venn diagram in **Fig. S8B**. Color bar shows average log_2_ fold change (in bold are genes directly implicated in immune escape mechanisms; underlined are genes actively involved in inflammatory processes). (**B**) Dot plots depicting percentages of CD163^+^ cells among CD14^+^CD16^-^ classical monocytes in HDs and GBM patients determined by multicolor flow cytometry analysis. Unpaired t-test (n=5 HDs, n=10 GBM patients (p-value: *: < 0.05). (**C**) Dot plot showing activated (red) or inactivated (blue) regulons comparing GBM patients (n=10) and HDs (n=5) (adjusted p-value < 0.05, |Avg_Log2FC| ≥ 0.25) across monocytic subsets (shared inactivated regulons are highlighted). Circle diameter denotes difference score between GBM patients and HDs.

Overall, the identification of these specific monocytic subsets was obtained through batch corrected analysis of re-clustered monocytes (**Fig. 4A-B**). However, we observed critical monocyte-related interpatient variation by CyTOF (**Fig. 2D**) and scRNA-seq analyses (**Fig. 3D-E**). Therefore, we re-clustered monocytic cells without correction to preserve patient-specific differences although considering the limitations imposed by the small cohort. We detected 20 clusters, clearly discriminating GBM patients and HDs as well as classical and non-classical monocytes, with classical monocyte clusters being individual-specific (**Fig. S9A-F**). Subsequently, we analyzed the expression levels of the 35 genes characterizing the identified GBM-CMS across patients and observed prominent specificities, especially for patients “GBM8” and “GBM10”, respectively clusters 3 and 9, showing a diverse signature compared to the others (**Fig. S9G**). Similarly to the other patients, they exhibited downregulation of MHC class II markers, including *HLA-DRB5*, but no upregulation of inflammation-limiting mediators, such as *IL1R2* and *CD163* (**Fig. S9G**). Importantly, these differences were not linked to dexamethasone treatment (GBM8 Dex+ - GBM10 Dex-) (**Fig. S9H**) or by specific variations in clinical parameters, such as genetic aberrations (**Table S1**), indicating unrelated inter-patient heterogeneity. Additionally, the segregation of patients based on dexamethasone therapy (Dex-, n=3; Dex+, n=6) did not lead to expression changes of CD11c and HLA-DR markers (**Fig. S9I**), indicating that the downregulation of antigen presenting cell markers is attributable to GBM and not simply to glucocorticoid treatment.

Taken together, GBM-CMS is defined by impaired MHC class II antigen presentation and increased expression of inflammation-controlling genes, with variability observed among individual patients.

### Non-classical monocytes in GBM patients display impaired inflammatory, antigen presentation and lineage programs

Following the characterization of the main classical monocytic populations, we sought to investigate if similar signatures were also detectable in non-classical monocytes. To do so, we analyzed the transcriptional differences between GBM patients and HDs within the corresponding cluster (**Fig. 7A**) and detected 249 DEGs (adjusted p-value < 0.05, |Avg_log_2_FC| ≥ 0.25) (**Table S12**). Non-classical monocytes in GBM patients were characterized by the up-regulation of complement-related genes (e.g., *C1QA*, *C1QB*, *C1QC*) along with down-regulation of inflammatory and MHC class II markers (e.g. *IL1β* and *HLA-DRB5*) (**Fig. 7B**). Non-classical monocytes shared 26 DEGs (approx. 75%) with GBM-CMS, including overexpression of *CD163, FKBP5* and *ZBTB16* (**Fig. 7B-C**). In line with these mRNA results, intermediate and non-classical monocytes in GBM patients showed lower expression levels of HLA-DR and CD11c compared with corresponding healthy cells also at the protein level (**Fig. 7D**, **Fig. S5C**). Further, they displayed increased expression of genes related to “cytoplasmic translation”, “regulation of immune effector process” and “macrophage activation” together with reduction of prototypical features, such as “leukocyte migration”, “chemotaxis” and “positive regulation of cell adhesion” (**Fig. 7E**) (**Table S13**). Like APC classical monocytes, they showed increased activity of TFs involved in the polarization of myeloid populations toward immunosuppressive phenotypes, such as CEBPD (*33*). Additionally, we detected decreased activity of TFs highly expressed in non-classical monocytes, including PLAGL2, TCF7L2 and NR4A1 (*34-36*), together with TFs involved in monocytic differentiation towards macrophages or dendritic cells, such as RUNX3 and BATF3 (*37*) (**Fig. 7F-G**), indicating an effect of GBM on the phenotypic acquisition of non-classical monocytes, both related to their lineage specifications and differentiation ability.

**Figure 7.**
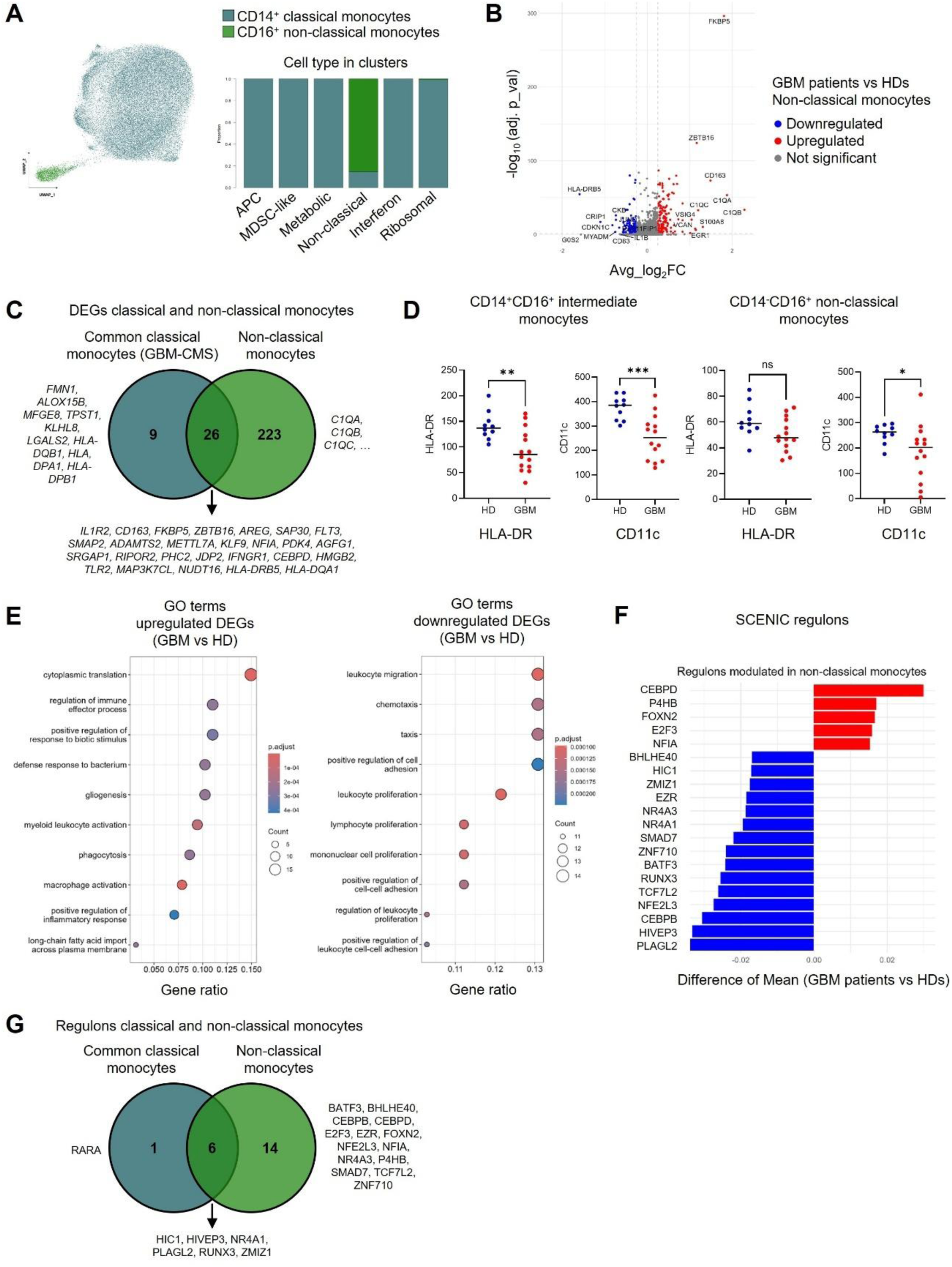
Non-classical monocytes in GBM patients display impaired inflammatory, antigen presentation and lineage programs. (**A**) UMAP and bar chart showing distribution of classical and non-classical monocytes in clusters formed by cells of HDs (n=5) and GBM patients (n=10). (**B**) Volcano plot showing top DEGs in non-classical monocytes comparing GBM patients with HDs (adjusted p-value < 0.05, |Avg_log2FC| ≥ 0.25; **Table S12**). Up-regulated genes in red and down-regulated genes in blue. (**C**) Venn diagram showing shared and discrete DEGs between the classical monocyte signature (GBM-CMS) and non-classical monocytes. (**D**) Dot plot displaying the levels of expression of HLA-DR and CD11c in CD14^+^CD16^+^ intermediate and CD14^-^CD16^+^ non-classical monocytes extracted from GBM patients and HDs determined by mass cytometry. Unpaired t-test (n=14 GBM patients, n=10 HDs; p-values: *: < 0.05; **: < 0.01; ***: < 0.001; ns: not significant). (**E**) Dot plots depicting main GO biological processes (BP) associated with up-regulated (left) and down-regulated (right) genes comparing GBM patients and HDs in non-classical monocytes (**Table S13**). Color bar denotes adjusted p-value and circle diameter depicts gene count of modulated genes within the GO-BP term. (**F**) Bar chart showing activated (red) and inactivated (blue) regulons in non-classical monocytes comparing GBM patients and HDs. (**G**) Venn diagram showing shared and distinct regulons between classical monocyte common TFs and non-classical monocytes.

In summary, GBM progression disrupts non-classical monocyte development, impairing their inflammatory and antigen-presenting functions, as in classical monocytes, along with lineage commitment and macrophage differentiation.

### Peripheral blood lymphocytes in GBM patients show transcriptional traits indicative of specific immune activation states

To further investigate how GBM progression affects adaptive immunity, we examined T cells, B cells and NK cells (**Fig. 3A**). In line with our method for analyzing monocytic cells, we re-clustered these subsets. We considered 18 non-overlapping clusters (**Fig. 8A**), identified using the same approach as in the myeloid cell analyses. We assigned four clusters to CD4^+^ T cells (clusters 1-4), three clusters to CD8^+^ T cells (clusters 5-7), two clusters to NK cells (NK CD56^dim^ in clusters 8, CD56^bright^ in cluster 9) and three clusters to B cells (cluster 10, B-naive in cluster 11 and B-intermediate in cluster 12). Moreover, six small clusters were composed of MAIT cells (cluster 13), CD56^+^ T cells (cluster 14) platelets (cluster 15), proliferating T cells (cluster 16), erythrocytes (cluster 17) and plasmablasts (cluster 18). We confirmed the identities of those clusters by analyzing the expression levels of corresponding cell-type specific markers (**Fig. 8B**). Following the grouping of the previous clusters into CD4^+^ T cells, CD8^+^ T cells, other T cells, NK cells and B cells, we quantitatively confirmed the cluster composition by cell type, delineating the identities of the main groups (**Fig. S10A**). Similarly to monocytes, PCA of these distinguished lymphocytes demonstrated that transcriptional weighted average proportion variance was primarily driven by cell subtypes (weighted average: 0.48) with minimal contribution from sample, batch, group and dexamethasone treatment (**Fig. S10B**). Both whole-cell gathering (**Fig. 3D**) and re-clustering of lymphocytic cells (**Fig. 8C**) showed only subtle transcriptional differences between lymphocytes derived from GBM patients and HDs, reflected also by the limited number of identified DEGs (adjusted p-value < 0.05, |Avg_log_2_FC| ≥ 0.25; **Table S14**). Collectively, clusters corresponding to CD4^+^ T cells (clusters 1-4), CD8^+^ T cells (clusters 5-7), NK cells (clusters 8-9) and B cells (cluster 12) in GBM patients showed features of lymphocytic cell activation, exemplified by shared prominent up-regulation of alarmin genes (e.g. *S100A8* and *S100A9*), which act as danger signals that trigger and modulate immune responses possibly contributing to the accumulation of MDSCs, along with down-regulation of genes associated with mitochondrial metabolism (e.g. *MT-CO2, MT-ATP6*) and AP-1 family (e.g. *JUNB, JUND*) (**Fig. 8D**), whose alterations are linked to anergic lymphocytes (*38*). Classical T cell exhaustion markers, such as TIGIT, PD-1, TIM-3 and LAG-3, were not modified at the transcriptional level. At the protein level, we did not detect increased expression levels of PD-1 and TIGIT in GBM patients, neither in CD4^+^ T and CD8^+^ T cells nor in NK CD56^dim^ and CD56^bright^ cells (**Fig. S10C**). However, genes linked to T cell function (e.g. *STAT1-3, JAK3, SOCS1-3*), trafficking (e.g. *CXCR4, CX3CR1*) and activation (e.g. *TNFAIP3, NFKBIZ*) were changed across CD4^+^ T cell clusters. Additionally, CD8^+^ T cells, particularly those in clusters 5-6, displayed reduced expression of HLA genes encoding MHC class II components (e.g. *HLA-DPA1, HLA-DQA2*), suggesting an impaired autocrine and paracrine activation capacity, with cluster 5 also characterized by decreased levels of the activation marker *CD69* (**Fig. 8D**).

**Figure 8.**
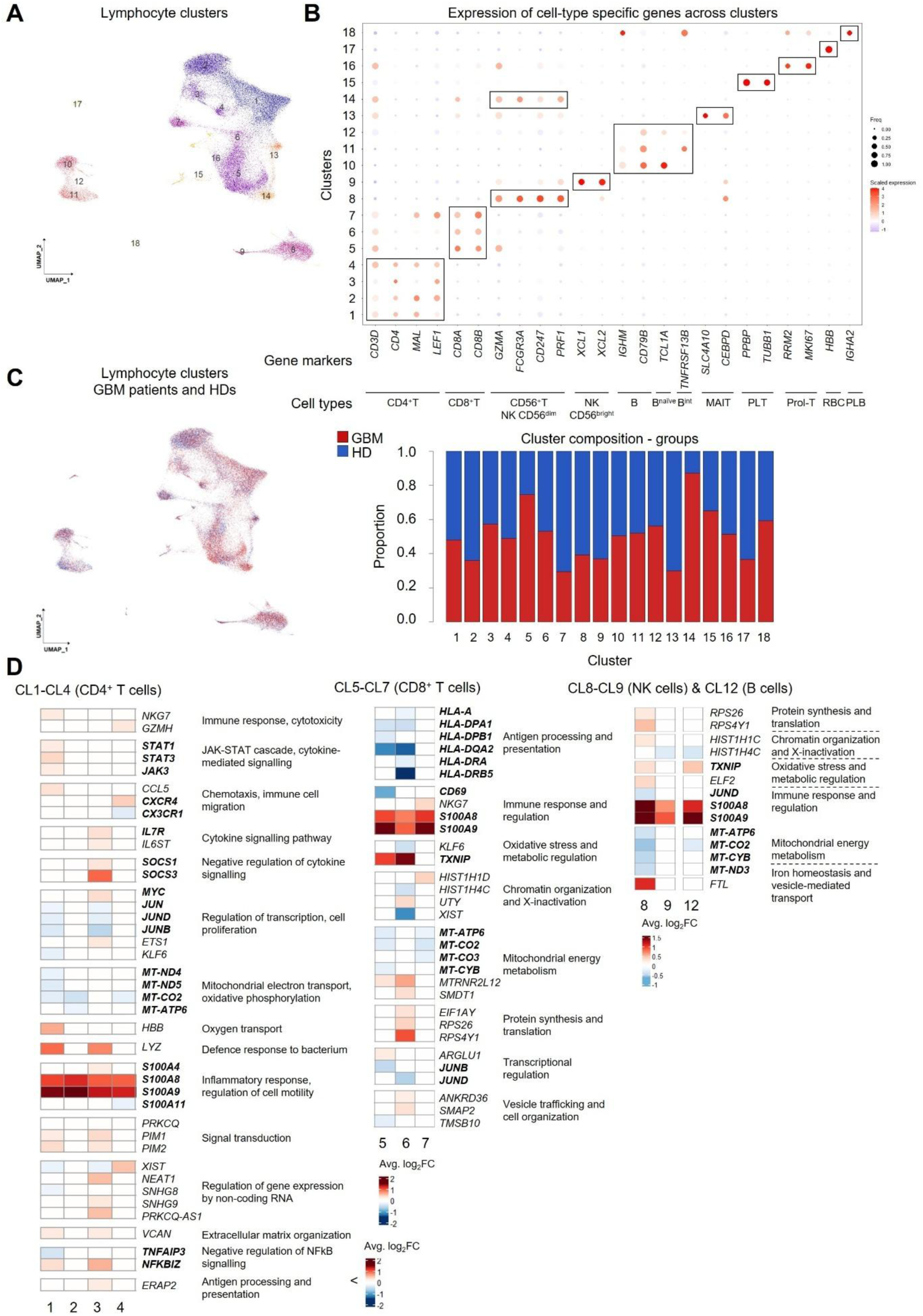
Peripheral blood lymphocytes in GBM patients show traits of immune dysfunction and exhaustion. (**A**) UMAP showing 18 clusters of lymphoid cells based on their transcriptional profiles. (**B**) Dot plot showing average expression of cell type markers in each cluster (adjusted p-value < 0.05, Avg_Log_2_FC ≥ 0.25). B^int^: intermediate B cells; MAIT: mucosal-associated invariant T cells; PLT: platelets; Prol-T: proliferative T cells; RBC: red blood cells; PLB: plasmablasts. (**C**) Left: UMAP depicting lymphoid cells from GBM patients (red) and HDs (blue). Right: Bar chart showing the proportion of cells derived from GBM patients and HDs across the 18 clusters. (**D**) Heatmap relative to DEGs (adjusted p-value < 0.05, |Avg_log_2_FC| ≥ 0.25; **Table S14**) detected in CD4^+^ T cells (CL1-CL4; left), CD8^+^ T cells (CL5-CL7; middle), NK cells (CL8-CL9; right) and B cells (CL12; right). Number of DEGs was minimal or not detected in the other clusters. Color bars show average log_2_ fold change (in bold genes associated with lymphocytic cell dysfunction and exhaustion).

T cells, NK cells and B cells exhibit shared upregulation of alarmin genes and downregulation of activation, mitochondrial respiration and metabolic genes, transcriptional hallmarks of lymphocyte dysfunction in GBM driven by intercellular or plasma factor perturbations.

### Impaired cell-cell crosstalk supports immune-suppressive trajectories in GBM patients

To assess if monocytic and lymphocytic changes in GBM arise from impaired intercellular communication or circulating mediators, we inferred PBMC ligand-receptor interactions using the OmniPath database (*39*) and performed differential interaction analysis with CellChat (*40*). This identified a prominent disruption of the interactome in patients (**Fig. 9A**). Specifically, we found monocytes (CD14^+^ and CD16^+^ subsets) and CD8^+^ T cells in HDs as having the highest outgoing and incoming interactions, respectively, indicating effective communication between innate and adaptive immune systems under homeostatic conditions. In GBM patients, we predicted prominent decreased outgoing and incoming interaction strengths across all main cell types, with CD8^+^ and CD4^+^ T cells being less affected compared to other cell types (**Fig. 9B**). These impairments were exemplified by changes in various signaling patterns, including decreased MHC-II outgoing signaling from both CD14^+^ and CD16^+^ monocytes, and reduced MHC-II incoming pattern in CD14^+^ cells (**Fig. 9C, Table S15**). As prominent ligand-receptor disruption underlying these observations, we identified HLA ligands (e.g. HLA-DRB5, HLA-DPA1, HLA-DPB1, HLA-DQB1 and HLA-DMA) expressed by CD14^+^ and CD16^+^ monocytes interacting with CD4 expressing CD14^+^ / CD16^+^ monocytes and CD4^+^ T cells (**Fig. 9D, Table S16**). Multicolor flow cytometry analyses of PBMCs further confirmed the decrease of HLA-DR and CD4 expression in GBM patient monocytes (**Fig. S5**). In naive / intermediate B cells, disruption of HLA ligands mainly impacted the putative interaction with CD4 expressing CD14^+^ cells, although the interaction between B cells and CD14^+^ monocytes is normally not heavily reliant on MHC-peptide presentation. Additional reduced ligand-receptor interactions were associated with cell-cell adhesion and stability of the immunological synapse, such as SELPLG - SELL and ICAM1 - SPN, which disruptions result in less effective T cell activation and antigen-specific immune responses, or linked to extravasation, such as IL16 - CD4 and the hemophilic binding CD99 - CD99 (**Fig. 9D, Table S16**). Overall, these observations were independent from dexamethasone treatment, both for interactions outgoing from myeloid (**Fig. S11A**) and lymphoid (**Fig. S11B**) populations.

**Figure 9.**
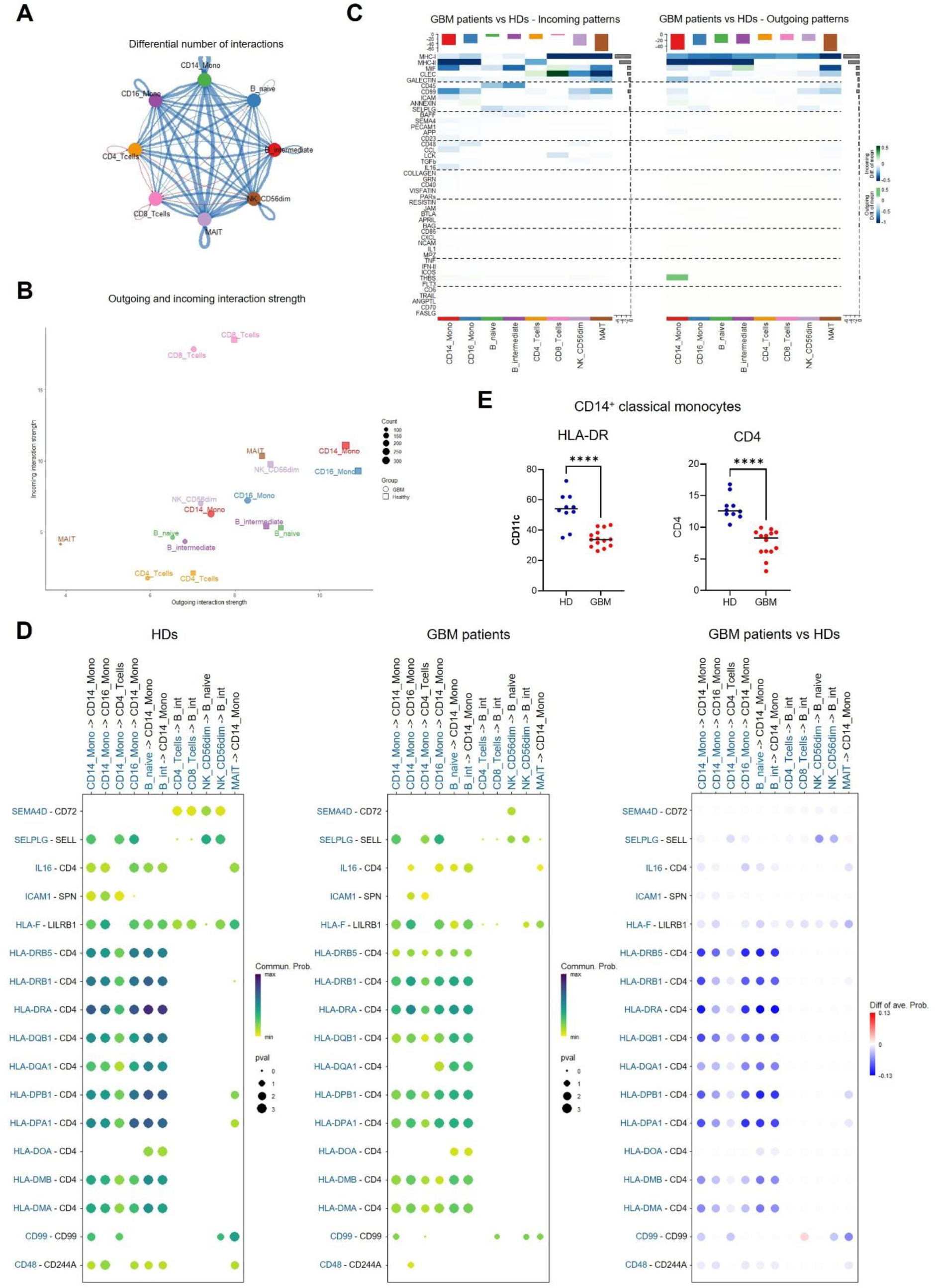
GBM patients display a disrupted cell-cell interactome in circulation. (**A**) Circus plot representing differential cell-cell communication networks between GBM patients (n=6) and HDs (n=3). Blue and red lines depict decreased or enhanced interaction, respectively, and line thickness represents number of interactions. (**B**) Scatter plot depicting outgoing (X-axis) and incoming (Y-axis) interaction strength per cell type showing average of HDs (squares; n=3) and GBM patients (circles; n=6). Square size or circle diameter represents number of counts. (**C**) Heatmaps showing average outgoing (left) and incoming (right) signaling patterns comparing GBM patients (n=6) with HDs (n=3). Rows and columns represent signaling patterns and cell types, respectively (**Table S15**). (**D**) Heatmaps visualizing a selected panel of ligand-receptor interactions in PBMCs from HDs (n=3; left), GBM patients (n=6; center) and differential regulation between the two groups (right). The dot color and size represent the calculated communication probability and p-values of differential communication, respectively. Significantly differentially regulated ligand-receptor pairs were calculated via the Wilcoxon rank-sum test. The first four columns represent interactions initiated by monocytic cells, whereas the remaining seven columns represent those started by lymphocytic cells (sender cells and their associated molecules in blue (**Table S16**). (**E**) Dot plot displaying the levels of expression of HLA-DR and CD4 in classical monocytes extracted from GBM patients and HDs determined by mass cytometry. Unpaired t-test (n=14 GBM patients, n=10 HDs; ****: p-value < 0.0001).

Collectively, all major circulating cell types in GBM patients show impaired cell-cell interactions, with CD14^+^ and CD16^+^ monocytes most affected. CD8^+^ and CD4^+^ T cells are less impacted, implicating soluble plasma factors over defective communication in their transcriptional changes.

## DISCUSSION

In our study, we found monocytic cells to be the most impacted immune cells during GBM development compared with lymphocyte populations, showing marked changes in their ratios and prominent transcriptional changes along with impaired lineage commitment. Specifically, we identified a GBM-classical monocytic signature (GBM-CMS) shared across the identified subsets characterized by the downregulation of MHC class II molecules and up-regulation of genes associated with immune evasion mechanisms, such as the decoy receptor *IL1R2* and the scavenger receptor *CD163*. Both receptors play important roles in restraining pro-inflammatory responses and are found on the cell surface or can be released into plasma as soluble factors (*41, 42*). Increased levels of HLA-DR^-^ MDSCs contribute to peripheral immune suppression and have been associated with worse prognosis (*13, 17, 20*). We confirmed the elevated abundance of MDSCs in the blood of GBM patients. High-grade glioma patient-derived CD14^+^HLA-DR^-^ monocytes can be precursors of MDSCs, also defined as licensing monocytes (*43*). Our trajectory analyses showed that MDSCs do not represent a terminal differentiation state, but rather a transitional phenotype within the monocytic compartment continuum. They originate from APC monocytes and subsequently give rise to metabolic program-enriched monocytic subsets possibly skewed by specific niches and cues they come across (*44*). The shared immature phenotype across the myeloid compartment suggests the existence of a dynamic bidirectional differentiation backbone in GBM patients, which acts as an immunosuppressive cell reservoir, continuously supplying the APC-like pool prone to differentiate into TAMs, while maintaining circulating APC monocytes in a hypofunctional state. This process is reminiscent of monocyte mobilization from bone marrow into the bloodstream to form a local reservoir in the spleen, which can be subsequently re-mobilized in response to injury or inflammation to differentiate into pro-resolving macrophages (*45, 46*). In this context, GBM patient monocytes may exhibit peripheral pre-skewing toward immunosuppression, facilitating Mo-TAM polarization in the TME. *In vitro* studies could confirm their greater propensity to form immunosuppressive subsets under GBM-like conditions versus HDs, identifying targets to block peripheral priming, recruitment and TME subversion while restoring anti-tumor immunity. Parallel studies assessing alterations in myeloid immune proportions and states into the bone marrow and spleen of GBM patients are warranted.

Patrolling non-classical monocytes, phagocytic cells with antigen presentation capacity and pro-inflammatory cytokine production under normal conditions (*47*), were reduced in GBM patients. Interestingly, a similar decrease has been described in patients affected by familial tauopathies (*48*) as well as in patients with Parkinson’s disease or amyotrophic lateral sclerosis (*49, 50*), thus reflecting a shared unknown mechanism associated with neurological insults. In line with previous studies showing that non-classical monocytes are absent in the TME (*51*) and, unlike classical monocytes, do not differentiate into Mo-TAMs (*52*), we identify APC monocytes as the main cell subset prone to give rise to Mo-TAMs in the TME. Targeting this monocytic subset to prevent tumor infiltration may be more effective than broadly modulating monocytes, which triggers compensatory neutrophil influx promoting GBM proneural-to-mesenchymal transition (*53*).

Lymphocytes in GBM did not show increased expression of typical exhaustion markers, such as TIGIT and PD-1. Instead, we observed down-regulation of mitochondrial genes critical for the electron transport chain and oxidative phosphorylation, suggesting these cells may have insufficient energy to perform their functions, such as proliferation and cytokine production, rendering them anergic (*54*). Suppressing mitochondrial activity in T cells, tumors can evade the anti-tumor effects of PD-1 blockade therapy (*55*).

Regarding the impact of GBM on cell-cell communication, the largely unchanged ligand-receptor interactions inferred between monocytic cells and both CD4^+^ and CD8^+^ T cells suggest that lymphocytic alterations are more likely driven by soluble factors, including secreted monocytic mediators favoring immune escape mechanisms, such as IL1R2, CD163, amphiregulin (AREG) and lactadherin (MFGE8), which genes exhibited prominent increased expression in our defined GBM-CMS. Further, increased alarmin expression, especially *S100A8/A9*, detected across most of our lymphocytic populations, can create a sustained inflammatory and tolerogenic environment that downregulates lymphocyte activation, proliferation, and cytokine function. These effects arise both directly via T cell receptor-mediated stress signaling and indirectly via myeloid cell reprogramming towards MDSCs and cytokine milieu alterations (*56*). Consequently, chronic alarmin elevation can constitute a major contributor to systemic lymphocytic dysfunction in GBM. Indeed, serum-derived soluble factors have been identified as key mediators of the widespread immunosuppressive state in GBM. Intriguingly, such systemic immune alterations, including peripheral lymphopenia, induced by serum-derived factors that are generally large, nonsteroidal molecules, are not specific to GBM. Similar changes have been reported in other CNS injuries (*57*), including stroke and viral encephalitis, and normally resolve once the underlying insult subsides (*9*). These systemic immune alterations point to a shared mechanism of CNS injury-induced immunosuppression across neuropathological conditions, creating challenges for specificity but also opportunities to extend insights into other brain diseases.

Among the limitations of the present study, while modest cohort sizes and reliance on pseudotime and ligand-receptor models respectively limit power and provide correlative rather than functional evidence, these can be addressed through orthogonal validation, including blocking assays, cytokine measurements, and cellular indexing of transcriptomes and epitopes by sequencing (CITE-seq). Clinically, these findings offer promising avenues to improve patient stratification and therapy guidance, particularly for immunotherapy, as well as to assess novel targets for anti-tumor activities. Key next steps include validating signatures in pre-diagnostic and treatment samples, integrating causal frameworks and functional assays, demonstrating added value over current diagnostics, and streamlining assays into practical formats like flow cytometry or qPCR/ELISA. Prospective studies will be essential to confirm impact and reproducibility.

In conclusion, understanding the cellular and molecular mechanisms driving the heterogeneous phenotypic changes of peripheral innate and adaptive immune cells may ultimately enable their reprogramming into fully competent immune effectors, offering a complementary and innovative avenue to restore and enhance anti-tumor immunity beyond current immune-stimulatory strategies (*58*). Notably, CAR-monocytes may be considered for these perspectives. Compared to macrophages, monocytes are easier to isolate, require no complex *in vitro* differentiation and exhibit superior tumor trafficking, making them a promising vehicle for targeting solid tumors, such as GBM (*59*). Therefore, our work not only provides valuable insights into novel diagnostic and therapeutic targets but also underscores the importance of understanding complex systemic immune cell dynamics for improving immunotherapeutic outcomes.

## MATERIALS AND METHODS

### Glioblastoma patients and healthy donors

Blood samples from GBM patients were collected at the Neurosurgical Department of the Centre Hospitalier de Luxembourg (CHL) as part of the PRECISION-PDX brain tumor collection from patients having given informed consent and with approval from the National Committee for Ethics in Research (CNER, AVIS 201201/06) in Luxembourg and according to the Declaration of Helsinki. Patient clinical data are summarized in **Table S1**. None of the patients underwent chemotherapy or radiotherapy prior to blood withdrawal. O6-methylguanine-DNA methyltransferase (MGMT) promoter methylation status testing and molecular GBM classification based on DNA methylation profiles were assessed via Illumina Infinium Methylation-Epic Array (Illumina, San Diego, CA, USA). GBM patients were considered dexamethasone-untreated if the treatment was not included in the patient management plan. Blood samples from HDs were collected at the Luxembourg Institute of Health within the HEALTHY BLOOD collection from individuals having given informed consent and with CNER approval (AVIS 202202/02).

### Isolation of mononuclear cells from whole blood

Whole blood was collected in BD Vacutainer ethylenediaminetetraacetic acid (EDTA) blood collection tubes (BD Bioscience, Franklin Lakes, NJ, USA) and directly processed for the isolation of PBMCs using SepMate™ tubes and Lymphoprep™ density gradient medium (STEMCELL Technologies, Vancouver, Canada), according to the manufacturer’s protocol. Briefly, the blood was first diluted 1:1 in phosphate buffered saline (PBS, Carl Roth, Karlsruhe, Germany) supplemented with 2% v/v fetal bovine serum (FBS, Gibco, Thermo Fisher Scientific, Waltham, MA, USA). The insert of SepMate™ 50 mL tubes was filled with 15 mL of Lymphoprep™ followed by the addition of diluted blood to form two distinct layers. After centrifugation (1200 x g for 10 minutes), the liquid fraction was collected and further washed twice with PBS supplemented with 2% v/v FBS. PBMCs were manually counted using a Neubauer chamber after 1:1 dilution in trypan blue (Thermo Fisher Scientific, Waltham, MA, USA) and cryopreserved at the concentration of 6-8 x 10^6^ cells/mL in Recovery™ Cell Culture Freezing Medium (Gibco, Thermo Fisher Scientific, Waltham, MA, USA) until subsequent analysis.

### Mass cytometry analysis

#### - Sample processing and analysis

Cryopreserved PBMCs from GBM patients and HDs were rapidly thawed at 37°C and diluted in pre-warmed RPMI 1640 medium (Gibco, Thermo Fisher Scientific, Waltham, MA, USA) complemented with 10% v/v FBS and 1% v/v penicillin/streptomycin (Lonza, Thermo Fisher Scientific, Waltham, MA, USA). After centrifugation (300 x g for 10 minutes), PBMC pellet was resuspended in 1 mL complete RPMI medium supplemented with 25 UI/mL benzonase (Merck Millipore, Darmstadt, German) and incubated at 37°C for 10 minutes to remove free nucleic acid contaminants. Following incubation, 3 x 10^6^ PBMCs were centrifuged (300 x g for 10 minutes), resuspended in PBS at a concentration of 10^7^ cells/mL and stained with 1 µM solution of the viability die Cell-ID Cisplatin (Standard BioTools, South San Francisco, CA, USA) for 5 minutes. After one wash with PBS supplemented with 5% FBS, cells were centrifuged (500 x g for 5 minutes) and resuspended in 270 µL Maxpar® Cell Staining Buffer (CSB, Standard BioTools, South San Francisco, CA, USA). To prevent unspecific staining, 5 µL of Human TruStain FcX Fc Receptor Blocking Solution (BioLegend, San Diego, CA, USA) was added, followed by a 10-minute incubation at room temperature. Surface marker staining was performed using the antibodies listed in **Table S17** diluted in CSB in a total volume of 300 µL. After 30 minutes of incubation at room temperature, cells were washed once and fixed with 1 mL of 1.6% v/v formaldehyde solution (Fisher Scientific, Waltham, MA, USA) for 15 minutes. Then, cells were incubated with 1 mL of 50 nM of Cell-ID Ir cationic nucleic acid intercalator solution (Standard BioTools, South San Francisco, CA, USA) overnight at +4°C. After one wash with CSB and one wash with Maxpar® Cell Acquisition Solution PLUS (CAS PLUS, Standard BioTools, South San Francisco, CA, USA), cell suspensions were filtered through a 35 µm cell strainer (VWR, Radnor, PA, USA), and counted with CASY counter (OLS, Bremen, Germany) according to the manufacturer’s instructions. Cells were then resuspended in CAS PLUS solution at a final concentration of 6 x 10^5^ cells/mL. Cells were acquired using a CyTOF XT mass cytometer (Fluidigm, South San Francisco, CA, USA). Data were normalized using the EQ™ Six Element Calibration Beads (Standard BioTools, South San Francisco, CA, USA). Data were analyzed with FlowJo software (10.8.1) (BD Bioscience, Franklin Lakes, NJ, USA).

#### - Unsupervised analysis

Data cleaning was performed by sequentially gating for singlet CD45^+^ live cells using FlowJo. Cleaned data were exported and uploaded to CellEngine (April 2025, CellCarta, Montreal, Quebec, https://cellcarta.com/cellenginesoftware/). Unsupervised clustering and dimensionality reduction were performed using the FlowSOM and t-SNE algorithm, respectively. FlowSOM and t-SNE were run simultaneously on arcsinh (x/5) transformed data subsampled to a maximum of 400,000 cells per sample to obtain 8,983,721 cells. Enriched FCS were exported in TSV format and imported in Tableau Desktop using the data preparation tool Tableau Prep as previously described (*60*).

#### - Cell subset classification based on surface marker expression

To enable cell subset assignment, baseline thresholds were defined for the median expression of each of the 33 surface markers to distinguish marker-positive from marker-negative cells. These thresholds were determined based on the overall distribution of marker expression across all metaclusters, pooled from all individuals in the cohort. Data visualization and plotting were performed using Tableau (Tableau Software, LLC, Seattle, WA, USA) as previously described (*60*). For each cluster, the threshold value of each marker expression was subtracted from its corresponding median expression. Values falling within the range -0.3 to 0.3 were classified as dim expressions. Clusters were subsequently classified based on their composite marker expression profiles using a sequential, gating-based strategy (**Table S2**).

#### - Statistical and trajectory analysis

CyTOF expression median values were Z-score normalized prior to downstream analysis to enable cross-marker comparison. Statistical testing was conducted using the Mann-Whitney U-test or Wilcoxon test, using Qlucore Omics Explorer (3.9, Qlucore, Lund, Sweden; https://qlucore.com/), GraphPad Prism (version 10.6.1 (892), GraphPad Software, LCC, San Diego, CA, USA; www.graphpad.com), or the *coin* package (version 1.4-3) in R (version 4.2.3). Adjusted p-values were calculated in Qlucore Omics Explorer using the Benjamini-Hochberg method to control false discovery rate (FDR) < 0.05.

Heat map and hierarchical clustering were visualized with *pheatmap* package (version 1.0.13), based on Z-score normalized median expression values of cell surface markers. Volcano plots were generated with *ggplot2* (version 3.5.2) to illustrate the relationship between fold changes and statistical significance of features identified by Mann-Whitney U-test (Qlucore). Significance thresholds were set at a q-value < 0.05. Color scales were chosen to enhance interpretability and emphasize relative differences across the data matrix. Labels were added to significant features using *ggrepel* (version 0.9.4).

Trajectory analysis was performed to investigate dynamic changes and progression patterns in monitored immune cell populations. Following data pre-processing, including dimensionality reduction and clustering, unsupervised K-means clustering with a resolution of 100 metaclusters was computed using CellEngine (April 2025, CellCarta, Montreal, Quebec, https://cellcarta.com/cellenginesoftware/).

Clusters were visualized by t-SNE using Tableau software (2025.1, Salesforce, San Francisco, CA, https://www.salesforce.com). To infer trajectories, pseudotime analysis was applied to macroclusters generated aggregating myeloid clusters, based on shared modulation trends observed for classical monocytes, MDSCs, and intermediate/non-classical monocytes, as described in the second paragraph of the Results section to order the macroclusters along putative phenotypic paths. A minimum spanning tree (MST) was built from the resulting pseudotime and trajectory coordinates, derived from principal component analysis (PCA) of Z-score normalized median expression values and visualized using *igraph* (version 1.5.1), *ggraph* (version 2.1.0), *tidygraph* (version 1.3.0) and *ggplot2* packages in R. Nodes were colored by pseudotime to reflect temporal progression.

To illustrate marker dynamics along inferred trajectories, directional arrows were overlaid, indicating the approximate timing and direction of involvement of the selected markers. Markers were annotated to highlight their contributions along the phenotypic paths (*grid* (version 4.2.3), *ggrepel* (version 0.9.4), *viridis* (version 0.6.5) and *dplyr* (version 1.1.4)).

PCA was performed with Qlucore Omics Explorer to assess heterogeneity across GBM patients and HDs. The contribution of each metaclusters to heterogeneity was profiled and overlaid to PCA via biplot analysis in R (*tidyverse* (version 2.0.0), *FactoMineR* (version 2.9) and *factoextra* (version 1.0.7)).

#### - Manual analysis

Cell subset assignment was carried out applying manual sequential gating strategy to CyTOF data with FlowJo software (10.8.1; BD Bioscience, Franklin Lakes, NJ, USA; **Table S5** and **Fig. S2**).

### Single-cell RNA sequencing analysis

For scRNA-seq analyses, PBMCs from four GBM patients (GBM1, GBM2, GBM3 and GBM4) and two HDs (HD1 and HD2) were pre-processed for CD14^+^ cell enrichment. Whole PBMCs from five GBM patients (GBM6, GBM7, GBM8, GBM9 and GBM10) and three HDs (HD3, HD4 and HD5) were directly processed. To assess the impact of CD14^+^ selection on the monocyte transcriptome, one patient sample (GBM5) was processed both as whole PBMCs and after CD14^+^ enrichment.

Cryopreserved PBMCs were rapidly thawed at 37°C and diluted in pre-warmed RPMI 1640 medium as mentioned above (see Mass cytometry analysis). After two washes with PBS supplemented with 0.05% v/v bovine serum albumin (BSA, Sigma-Aldrich, St. Louis, MO, USA) and EDTA 200 nM, one million PBMCs were incubated for 20 minutes with CD14 beads (130-118-906, Miltenyi Biotec, Bergisch Gladbach, Germany) and passed through LS columns (130-042-401, Miltenyi Biotec, Bergisch Gladbach, Germany) to enrich CD14^+^ cells by magnetic separation. Lastly, CD14^+^ cells or whole PBMCs were collected and further processed for scRNA-seq analyses. Cell suspension was filtered through a 50 µm filter (PluriSelect, Leipzig, Germany) to remove aggregates. Cell viability and concentrations were assessed via Vi-CELL automated cell counter (Beckman Coulter, Brea, CA, USA) and C-Chip Disposable Hemocytometer (NanoEntek, Seoul, South Korea). Cell suspension was subsequently diluted to obtain a final concentration of 1 x 10^4^ cells/µL following 10x Genomics guidelines (10x Genomics, Pleasanton, CA, USA). The single-cell suspension was loaded onto a Chromium Next GEM chip and processed using the Chromium Controller (10x Genomics, Pleasanton, CA, USA) to encapsulate single cells and Gel Beads-in-Emulsion (GEMs). The scRNA-seq libraries were prepared using the Chromium Next GEM Single Cell 3’ GEM, Library & Gel Bead Kit v3.1 (10x Genomics, Pleasanton, CA, USA), which includes reagents for reverse transcription, cDNA amplification, and library construction, and the Chromium i7 Multiplex Kit (10x Genomics) for sample barcoding. Libraries were purified using SPRIselect magnetic beads (Beckman Coulter) and quality assessed using an Agilent 2100 Bioanalyzer (Agilent Technologies, Santa Clara, CA, USA). Sequencing was performed on a NextSeq 500/550 system using the High Output Kit v2.5 (150 cycles; Illumina) or NovaSeq 6000 system using a S1 flow cell (Illumina), with an estimated depth of at least 30,000 reads per cell.

Raw sequencing data were processed using Cell Ranger v7.0.1 to generate gene-barcode UMI count for each sample with included introns option set to false. Outputs of Cell Ranger were analyzed in R/CRAN software (version 4.4.1; R Core Team 2024). Cell-calling analysis was performed to remove empty barcodes. For each sample, the inflection point in the plot of log(ranks) vs log(total counts) was calculated using the barcodeRanks function of *DropletUtils* R package (*18*). Barcodes with total UMI counts above that point were called cells. Cells from all samples were then united into a single Seurat object and analyzed in the Seurat v4 framework (*18*). First, barcodes with more than 20% of UMI counts originated from mitochondrial genes were removed. Gene expressions were then log-normalized and scaled using the NormalizeData (scale factor = 10,000) and the ScaleData functions. To reduce data dimensionality, PCA was run on scaled-normalized expression of the 5,000 most variable genes identified using the FindVariableFeatures function. PCA was limited to 200 principal components among which only the top 50 were input for the RunUMAP function for 2D-UMAP calculation. For clustering analysis, a shared nearest neighbor (SNN) graph was constructed from the 50 first PCs using the FindNeighbour method. Cells were then grouped according to their transcriptional similarities using FindClusters (resolution = 0.05). Cell type identification was performed using the standard marker-based scoring approach in conjunction with clustering analysis. Canonical gene markers of major PBMC cell types were obtained from the Human BioMolecular Atlas Program Data Portal (*18*). For each cell, the overexpression score of each cell type was estimated using the AddModuleScore function. Cell types were assigned according to the highest module score. For the whole PBMC dataset, Azimuth 2021 level 1 was used to discriminate the major cell types.

Cell-cell communication was inferred using the R package *CellChat* (v2.1.2) (*40, 61*). CellChat estimates intercellular signaling probabilities between cell populations based on prior knowledge of ligand-receptor (LR) interactions. CellChatDB (human, version 1) was used as reference database. Analysis was performed at sample level and restricted to the nine PBMC enriched samples. To minimize noise and ensure robust inference, only the major cell types identified in the single-cell dataset were included: CD14^+^ monocytes, CD16^+^ monocytes, CD4^+^ and CD8^+^ T cells, B-naive, B-intermediate and MAIT cells when sufficiently abundant. Each sample was pre-processed using the standard CellChat workflow. Briefly, the sample’s data was retrieved from the whole single-cell Seurat Object and subset to the cell types of interest. A CellChat object was then created based on the log-normalized expression data, grouping cells by their annotated cell type. Subsequently, the CellChat object was reduced to signaling genes using the subsetData, followed by identification of overexpressed genes and overexpressed interactions using the identifyOverExpressedGenes and identifyOverExpressedInteractions functions, respectively. Intercellular communication probabilities were estimated using the computeCommunProb function with the “triMean” method. Interactions supported by less than 10 cells per cell type were discarded. Communication probabilities were further aggregated at the pathway level to build global signaling networks using sequentially the computeCommunProbPathway and aggregateNet functions. Lastly, network centrality measures (in-degree, out-degree, betweenness, and closeness) were computed to identify major sender and receiver cell populations within each signaling pathway (netAnalysis_computeCentrality function, slot.name = "netP").

Visualization of sample-level CellChat results were performed using CellChat’s plotting functions or in-house R scripts built on *ggplot2* (version 3.5.2), *pheatmap* (version 1.0.13), *ComplexHeatmap* (*62*) (version 2.16.0) and R base graphics. For CellChat functions, custom adaptations were implemented to enable consistent comparison across groups of multiple samples.

To compare the strength of specific cell-cell communication axes between GBM patients and HDs, differential analysis based on the inferred communication probabilities was conducted. For each sample, LR probabilities were extracted from CellChat object. Weights were aggregated across all samples to construct a comparative matrix for each unique source-target-LR interaction. Interactions with a sum of weights equal to zero across all samples were excluded. For each specific LR pair within a given source-target link, an unpaired Student’s t-test was performed to compare the mean interaction weights between the two groups. Raw p-values were adjusted using the Benjamini-Hochberg correction and the mean differences were calculated to quantify the magnitude of change. Source-target-LRs were selected based on adjusted p-value < 0.05.

### Statistical analysis

Statistical analysis of mass cytometry data was performed using Mann-Whitney U-test followed by Benjamini-Hochberg correction. Statistical analysis of flow cytometry data was performed using t-test in GraphPad Prism (version 10.6.1 (892), GraphPad Software, LCC, San Diego, CA, USA; www.graphpad.com). To determine differentially expressed genes (DEGs) of the scRNA-seq dataset, Wilcoxon Rank Sum test, AUC and logistic regression were applied. Genes with |log_2_FC| ≥ 0.25 and adjusted p-value < 0.05 were considered as DEGs.

## Supporting information

Supplementary materials and methods, figures, and legends for data files

## List of Supplementary Materials

Materials and Methods

Figs. S1 to S11

Legends for data files S1 to S17

References

## Acknowledgements

LIH PRECISION-PDX brain tumor collection was supported by the Neurosurgery Department of the Centre Hospitalier de Luxembourg, the NORLUX Neuro-Oncology Laboratory and the Clinical and Epidemiological Investigation Centre of LIH. We are grateful to the NORLUX Neuro-Oncology Laboratory and Ms. Amandine Bernard for technical support with the isolation of PBMCs from GBM patients and HDs. We thank the LUXGEN platform for their support with single-cell sequencing. Schematic figures were created using BioRender.com. For the purpose of open access, and in fulfilment of the obligations arising from the FNR grant agreement, the author has applied a Creative Commons Attribution 4.0 International (CC BY 4.0) license to any Author Accepted Manuscript version arising from this submission.

## Funding

Fondation du Pélican de Mie et Pierre Hippert-Faber under the aegis of Fondation de Luxembourg (AnS) Luxembourg National Research Fund (FNR) through the FNR-PRIDE programs for doctoral education PRIDE21/16763386/CANBIO2 (EC)

Luxembourg National Research Fund (FNR) through the FNR-PRIDE programs for doctoral education PRIDE/14254520/I2TRON (FL-HM)

Luxembourg National Research Fund (FNR) CORE grant C21/BM/15739125/DIOMEDES (BN, PVN, AG)

Fonds de la Recherche Scientifique (FNRS)-Télévie grant GBModImm no. 7.8513.18/7651720F (EK, SPN, AG)

Fonds de la Recherche Scientifique (FNRS)-Télévie grant ImmoGBM 7.8505.20/7.6603.22 (EK, SPN, AG)

Luxembourg National Research Fund (FNR) grant INTER/DFG/17/11583046 (AlS)

Luxembourg National Research Fund (FNR) PEARL grant P16/BM/11192868 (MM)

Luxembourg National Research Fund (FNR) and Fondation Cancer CORE grant C24/BM/18858278/GRALL (AP, AM)

Personalised Medicine Consortium (PMC) of Luxembourg Pump Prime Fund (PIMP-GBM)

Action LIONS « Vaincre le Cancer » Luxembourg

Luxembourg Institute of Health (LIH)

## Author contributions

Conceptualization: AnS, AM

Methodology: AnS, TK, CC, KG, BN, AlS, AC, PVN, AG, AP, AM

Resources: FH, GB, MM, BM, SPN, PVN, AG, AM

Formal analysis: AnS, TK, CC, EC, BN, FL-HM, EK, RDC, AC, AP, AM

Data curation and visualization, AnS, TK, CC, AM

Funding acquisition: AM

Supervision: AlS, AC, PVN, AG, AP, AM

Writing – original draft: AnS, CC, AM

Writing – review & editing: all authors

## Data and materials availability

The single-cell mass cytometry data has been deposited at Mendeley Data: Cerella, Claudia; Cosma, Antonio; Michelucci, Alessandro (2025), “Circulating immune profiling reveals impaired monocyte phenotypes and trajectories driving immunosuppression in glioblastoma”, Mendeley Data, V1, doi: 10.17632/gth6pct574.1 https://data.mendeley.com/preview/gth6pct574?a=5ef7cf0e-9eab-4840-9985-5a61efe975e3

The single-cell RNA-sequencing data has been deposited at the European Genome-phenome Archive (EGA), which is hosted by the EBI and the CRG, under accession number EGASXXXXXXXXXXX (ongoing). Further information about EGA can be found on https://ega-archive.org "The European Genome-phenome Archive in 2021" (https://academic.oup.com/nar/advance-article/doi/10.1093/nar/gkab1059/6430505).

## Ethical approval and consent to participate

The collection of blood and tumor tissue from glioblastoma patients is a procedure routinely carried out in the framework of a collaboration between the Neurosurgery Centre at the Centre Hospitalier de Luxembourg (CHL) and the NORLUX Neuro-Oncology Laboratory at the Luxembourg Institute of Health (LIH) (www.precision-pdx.lu). All patients received a detailed information sheet and signed an informed consent form to agree to provide blood and tissue for research. The procedure was approved by the National Ethics Committee for Research (CNER, PRECISION-PDX, AVIS 201201/06), and the data collection including clinical and genetic data was authorized under pseudo-anonymised format and supervised by the National Data Protection Committee (CNPD, 929/2017) in Luxembourg. The CNER also approved the collection of blood samples from HDs (HEALTHY BLOOD, AVIS 202202/02).

