## Supplementary materials and methods, figures, and legends for data files for "Circulating immune profiling reveals impaired monocyte states and trajectories driving immunosuppression in glioblastoma"

#### Single-cell RNA-sequencing bioinformatics analysis

- Myeloid cells

Cells annotated as dendritic cells or monocytes from clusters 2 to 7 (48,814 cells) were re-analyzed separately. Raw UMI counts were log-normalized using a scale factor of 10,000 followed by scaling using the ScaleData function. To discriminate between subpopulations, marker-based module scores were calculated using AddModuleScore with the signatures of human PBMCs obtained from Azimuth 2021 level 2 as reference.

For dimensionality reduction, the 10,000 most variable genes identified using FindVariableFeatures were used to compute 200 principal components (PCs) with RunPCA. To evaluate the biological relevance of individual PCs, we quantified their association with technical and biological covariates (sample/patient, cell extraction protocol, gender, group, batch, cell type annotations and cell size). Each PC was regressed independently against each covariate using the lm function of the stats package (R Core Team, 2024), and the adjusted $R^2$ was extracted, which is an estimate of the variance attributable to that covariate. The resulting PC-covariate association matrix was visualized as a heatmap, enabling the selection of PCs minimally influenced by technical effects for UMAP. To complement this PC-level analysis, Principal Variance Component Analysis (PVCA) was performed to estimate the relative contribution of each factor to total transcriptional variance, considering multiple covariates simultaneously. Consistent with the PC regression analysis, PVCA revealed pronounced patient- and batch-associated effects. Given the magnitude of these effects, datasets were subsequently integrated using Sub-Type Anchor Correction for Alignment in Seurat (STACAS) to mitigate sample- or batch-driven variation. Monocyte Seurat Object was split by sample and Run.STACAS function of *STACAS* package [STACAS: Sub-Type Anchoring Correction for Alignment in Seurat. R package version 2.2.2] was used with cell subtypes as cell labels to guide the alignment. STACAS integrated data was then used for visualization and clustering. For visualization, STACAS was projected into a 2D space using RunUMAP function. Clustering analysis was performed with FindNeighbors followed by FindClusters across a range of resolutions (0.05-0.5, step 0.05) to explore different levels of granularity. A resolution of 0.05 was selected for downstream analysis, yielding a total of six clusters. For cluster annotation, differential expression analysis was first performed to identify cluster marker genes. Annotation of the six clusters was performed in two steps. First, differential expression analysis between clusters was carried out using Seurat's FindMarkers function of Seurat with MAST as statistical test (*1*) and sample included as latent variable to account for sample-specific effects. Cluster marker genes were defined as those with a $\log_2$ fold-change $\geq 0.25$ and FDR $< 0.05$. Second, the biological significance of these marker genes was obtained through over-representation analysis (see gene ontology described below).

The identified clusters, along with their associated markers, were compared to the genetic programs defined by Miller and colleagues. Similarity between gene lists was evaluated using the Jaccard index to assess the concordance and a binomial-based test to determine the significance. Overexpression of each program in the identified monocytic populations was calculated using AddModuleScore function, and the distribution of these scores across clusters was visualized with boxplots. To assess the usage of PBMC transcriptional programs defined by Miller and collaborators to confirm cell type identity of our monocyte subset, non-negative least square was used. Counts were extracted from Seurat objects and restricted to genes shared with the programs. Program contributions for each cell were estimated using nnls function of NNLS 1.6 R package (https://cran.r-project.org/web/packages/nnls/index.html). Resulting weights were normalized to percentages per cell and averaged across the identified six clusters. Cluster-level program enrichment was visualized as dot plots using ggplot2, with dot size and color reflecting mean program contribution.

To identify transcriptional changes between GBM patients and HDs, differential expression analysis was conducted either within each cluster or across all classical and non-classical monocytes. Differential expression was computed using Seurat's FindMarkers function with MAST as statistical test, sample as latent variable and group as grouping factor. Genes with $|\text{Log}_2$ fold-change$| \geq 0.25$ and FDR $< 0.05$ were considered differentially expressed.

Gene regulatory networks (GRNs) were inferred with the Single-Cell Regulatory Network Inference and Clustering (SCENIC) pipeline implemented in R (*SCENIC* package version 1.3.1). All databases were downloaded from the cisTarget databases (https://resources.aertslab.org/cistarget/). To start,

SCENIC was initialized with human gene annotations (hgnc) and the corresponding cis-regulation databases from RcisTarget package (version 1.16). The log-normalized gene expression count of all monocytes was extracted from the Seurat object. Genes were filtered using the SCENIC geneFiltering function, keeping only genes expressed in at least 1% of cells and with a minimum total count corresponding to 1% of the number of cells. Based on the resulting gene expression filtered matrix, a gene-gene correlation network was calculated using the runCorrelation function of *SCENIC*. Then, GRNs were inferred using the runGenie3 function with the filtered log-normalized gene expression matrix, all transcription factors available in the database, 20 CPUs, 200 data partitions and 500 trees per random forest. Following co-expression inference, gene regulatory modules were built with the runSCENIC_1_coexNetwork2modules and the runSCENIC_createRegulons functions. RunSCENIC_1_coexNetwork2modules help transform the raw output of runGenie3 into co-expression modules where transcription factors (TFs) are grouped with their potential target genes (TGs). The co-expression modules were then refined into biologically meaningful regulons using runSCENIC_createRegulons (coexMethods=c("top1sd","top50perTarget")), which allows only TF-TG supported by binding motifs in the regulatory region of the gene candidates. As a final step, runSCENIC_3_scoreCells was applied to quantify regulon activity at the single-cell level with aucThreshold = 0.2 and auxMaxRank = 500 to ensure detection of weaker regulons. For a given regulon, the activity score represents the proportion of its target genes highly expressed within a cell. These scores were subsequently used for clustering, dimension reduction and comparative analysis.

To determine transcriptional trajectories, we first ran the basic Seurat v5 pipeline (*2*) on the published Mo-TAM dataset by Yabo and colleagues and performed clustering using the res = 0.2 parameter of the FindClusters() function in Seurat, which resulted in seven clusters. Next, we integrated Mo-TAM dataset with our dataset by merging the two corresponding Seurat objects. The standard Seurat pipeline, including batch correction with Harmony (*3*), was run on the merged data as follows: NormalizeData > FindVariableFeatures > ScaleData > RunPCA > RunHarmony > RunUMAP (reduction = "harmony"). To account for differences in UMI counts, gene counts, and cell cycle states between the two datasets, we regressed out the variables nCount_RNA, nFeature_RNA, and Phase (cell cycle state) by passing them to the vars.to.regress parameter of the ScaleData function. Batch correction was performed by treating the Mo-TAM dataset as a single batch and using the sample column from the current dataset to define other batches. The cell clusters from both datasets before merging were kept unchanged after integration by labelling clusters from our dataset as 1-6 (as identified by the STACAS algorithm) and the seven Mo-TAM clusters as 7-13. These clusters were then provided to the TSCAN algorithm for cluster assignments for each cell, enabling trajectory analysis on the integrated data. For Minimum Spanning Tree (MST) and pseudotime calculations, we explicitly specified batch-corrected Harmony embedding in the TSCAN functions (e.g. use.dimred = "harmony").

- Lymphocytic cells

Lymphoid cells were extracted from the single-cell dataset based on the initial clustering (clusters 1 and 8-12) and further purified by removing cells that exhibited the highest overexpression score for a myeloid lineage signature. The resulting lymphocyte subset was then re-analyzed independently following the same Seurat pipeline used for monocytes. Briefly, data were log- normalized (scale factor = 10,000), scaled, and the 10,000 most variable genes were selected for dimensionality reduction. PCA (RunPCA, 200 PCs) was performed, followed by UMAP (RunUMAP, 50 first PCs) for visualization. For cell sub-type assignation, overexpression scores for the gene sets represent in the Azimuth Human PBMC level 2 (Azimuth 2021) were computed using addModuleScore. Each cell was assigned to the subtype with the highest score.

As for the monocytes, the contribution of each experimental variable to the overall variability and to each PC was assessed. Although cell type accounted for the largest faction of variance, patient and batch effects were still detectable. STACAS was therefore applied to mitigate inter-individual effects. The STACAS-corrected data were then subjected to UMAP visualization (RunUMAP, all PCs) and clustering (FindNeighbors, FindClusters with resolution = 0.25). The resulting 18 clusters were annotated manually by combining the cell sub-type generated using overexpression-based approach and the expression of prototypic, non-redundant cell type-specific genes. Differential expression analysis was conducted with the same procedures as for the monocytes.

**Flow cytometry analysis**

Thawed PBMCs were resuspended in PBS complemented with 0.2% v/v BSA (Sigma-Aldrich, St. Louis, MO, USA) and Human TruStain FcX Fc receptor Blocking Solution (BioLegend, San Diego, CA, USA) was used to avoid unspecific staining. Surface marker staining was performed using antibodies listed in **Table S17** diluted in BD Horizon™ Brilliant Stain Buffer (BD Bioscience, Franklin Lakes, NJ, USA). Cell suspensions were incubated with antibody mix for 30 minutes at +4°C. After two washes using PBS complemented with 0.2% v/v BSA, cells were incubated for 30 minutes at +4°C with 1 µL of Live/Dead fixable Near IR (Invitrogen, Carlsbad, CA, USA) in 100 µL PBS complemented with 0.2% v/v BSA. After two washes using PBS complemented with 0.2% v/v BSA, cells were fixed using BD Cytofix™ fixation buffer (BD Bioscience, Franklin Lakes, NJ, USA) for 20 minutes at +4°C. After two washes using PBS complemented with 0.2% v/v BSA, cells were resuspended in PBS complemented with 0.2% v/v BSA for acquisition. As negative control, cells were stained with Live/Dead fixable Near IR (Invitrogen, Carlsbad, CA, USA). BD™ CompBeads Anti-mouse Ig κ / Negative Control Compensation Particles Set were used for compensation. All the samples were recorded using NovoCyte Quanteon (Agilent Technologies, Santa Clara, CA, USA). The recorded data (FCS files) were analyzed with FlowJo software (10.8.1; BD Bioscience, Franklin Lakes, NJ, USA).

**Gene Ontology (GO) analysis**

Differentially expressed genes ($|\log_2 FC| \geq 0.25$, adjusted p-value $< 0.05$) were submitted to *clusterProfiler* R package (*4, 5*) (10.18129/B9.bioc.AnnotationDbi;10.18129/B9.bioc.org.Hs.eg.db) for Gene Ontology Biological Processes (GO_BP) enrichment analysis (GO_BP terms considered enriched with adjusted p-value < 0.05).

**Comparative quantification of myeloid macrocluster frequencies from CyTOF and myeloid cluster frequencies from single-cell RNA-sequencing data**

Cell frequencies expressed as percentages were calculated for each of the five myeloid macroclusters identified by CyTOF analysis. For each individual (10 HDs and 14 GBM patients), the total number of cells in each of the five macroclusters was obtained from the CyTOF recorded data and their frequency was calculated relative to the total $CD45^+CD66^-$ cell population. The proportion of cells assigned to each of the six myeloid clusters identified from the scRNA-seq analyses was obtained following STACAS analysis.

**Survival studies**

Survival analysis was generated using data from the Glioma Longitudinal AnalySis (GLASS) consortium (*6*) (https://glass-consortium.org). TPM gene expression values and clinical features were retrieved from cBioPortal (Diffuse Glioma (GLASS consortium); released version: May 31, 2022). The analysis included non-redundant patient samples, collected at the earliest follow-up (primary tumors; originally identified with the code TP) and profiled for *IDH* mutational status (n = 96 IDH-WT patients). Significantly differentially expressed genes ($log_2FC > 0.25$; adjusted p-value < 0.05) for each of the six monocytic clusters identified were used to generate cluster-specific gene signature scores. TPM gene expression values of significantly expressed genes across all six clusters were Z-scored and intersections with gene signature scores were analyzed in R (version 4.2.3). Patient samples were categorized according to the median gene score values and probability of overall survival was assessed by Kaplan-Meier method (R; *survival, survminer*, *ggsurvplot*).

**Ivy Glioblastoma Atlas Project dataset analysis**

We selected classical APC monocyte genes and Mo-TAM genes modulated across the monocytic differentiation into Mo-TAMs (**Table S11**) and took advantage of Single Cell Portal (https://singlecell.broadinstitute.org/single_cell) to exclude genes expressed by tumor cells. Next, a selection of classical APC monocyte genes and Mo-TAMs genes, together with microglia-specific genes (*SALL1, P2RY13, P2RY12* and *TMEM119*), were submitted to Ivy Glioblastoma Atlas Project to determine their expression in the following GBM areas: leading edge, infiltrating tumor, cellular tumor, necrotic tumor, pseudopalisade, hyperplastic blood vessels and microvascular proliferation.

### SUPPLEMENTARY FIGURES

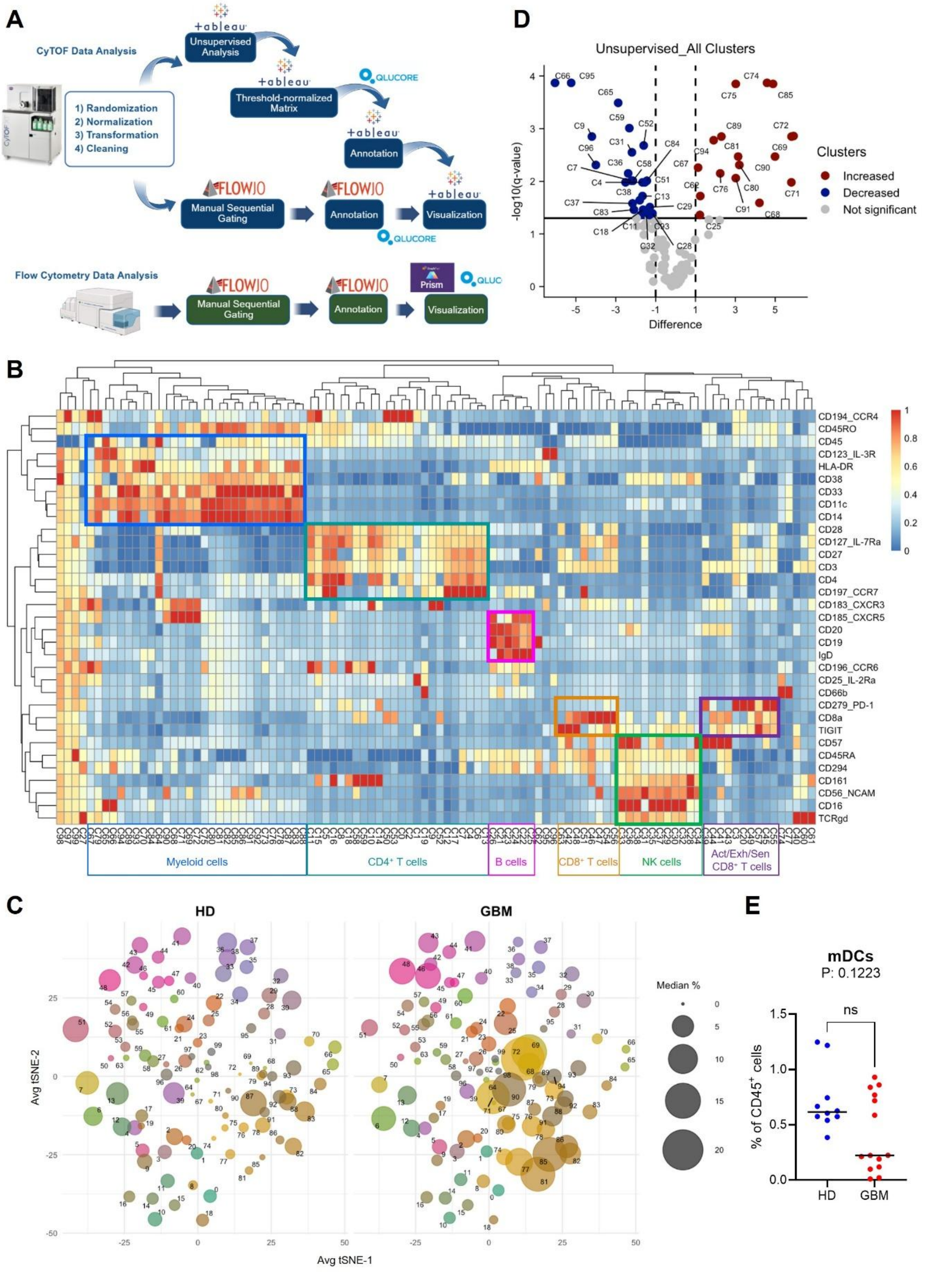


**Figure S1. Cell subset assignment from unsupervised analysis of the CyTOF dataset.** (**A**) Schematic showing profiling of circulating immune cells by mass and flow cytometry, using multiple software platforms for unsupervised and supervised analyses (created with BioRender.com). (**B**) Heatmap with hierarchical clustering of CyTOF data (HDs: n=10; GBM patients:

n=14) resolved in 100 metaclusters (unsupervised analysis). Colour bar denotes Z-scored values of mass intensity units. (**C**) Average t-SNE plots showing differential abundance of each metacluster between HDs and GBM patients. (**D**) Volcano plot of unsupervised clusters related to Mann-Whitney U test followed by Benjamini-Hochberg correction, computed with Qlucore on the same cohort of **Fig. 1**, prior to cell subset assignment. (**E**) Frequency of monocytic dendritic cells (mDCs) in HDs and GBM patients. Non-parametric two-sided Mann-Whitney U test (HD: n=5; GBM: n=10; ns: not significant).

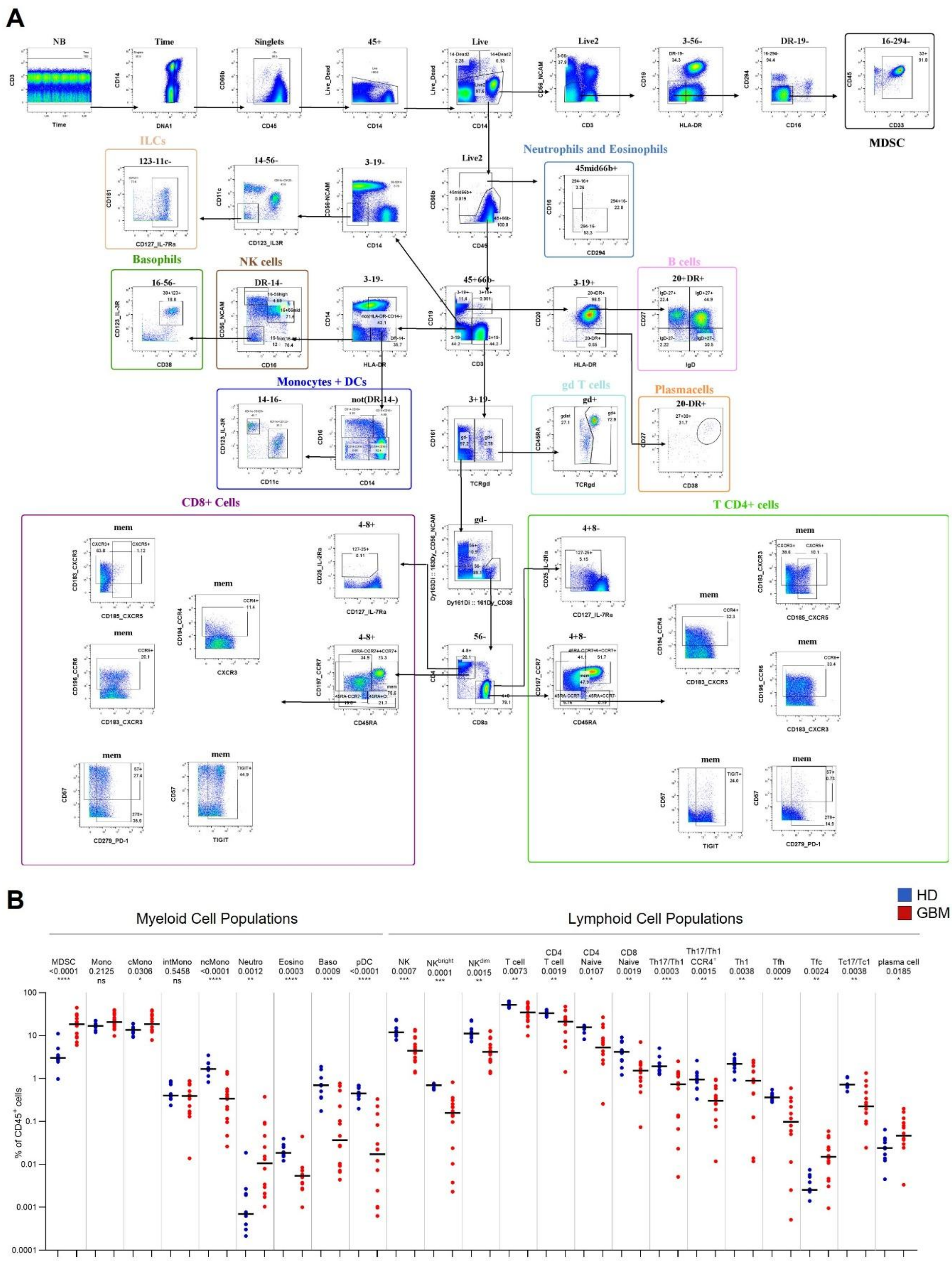


**Figure S2. Supervised analysis and cell subset assignment of CyTOF data.** (**A**) Sequential gating strategy used for supervised analysis (see also **Tables S4-S6**). (**B**) Distribution of significantly changed myeloid and lymphoid subsets in HDs (blue) and GBM patients (red). Individual data points are shown; median values are indicated by black lines. A logarithmic Y-axis was applied to visualize subsets together. Statistical analysis: non-parametric two-sided Mann-Whitney U test (p-values: *: < 0.05; **: < 0.01; ***: < 0.001; ****: < 0.0001; ns: not significant).

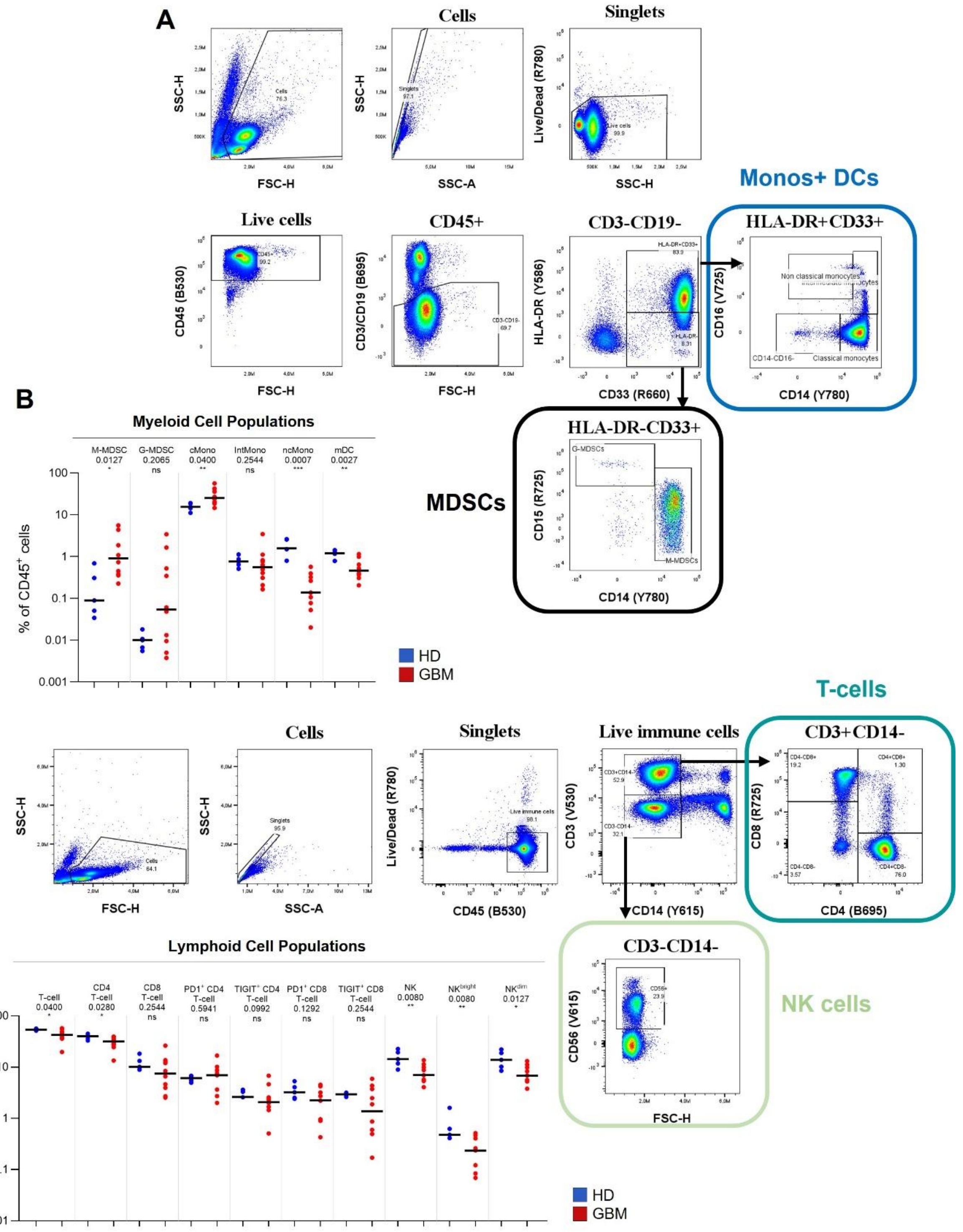


**Figure S3. Flow cytometry validation of cell subsets.** Sequential gating strategy applied to the cohort used in flow cytometry confirmatory studies (HDs: n=5; GBM patients: n=10) to identify (**A**) monocytes (Monos), mDCs and MDSCs or (**C**) T and NK cells. Cell subsets of (**B**) myeloid and (**D**) lymphoid compartments modulated between HDs (blue) and GBM patients (red) showing distribution at individual level. Median values are indicated by the black line. A logarithmic scale was applied to the

Y-axis to plot all different subsets together. Statistical analysis: non-parametric two-sided Mann-Whitney U test (p-values: *: < 0.05; **: < 0.01; ***: < 0.001; ns: not significant).

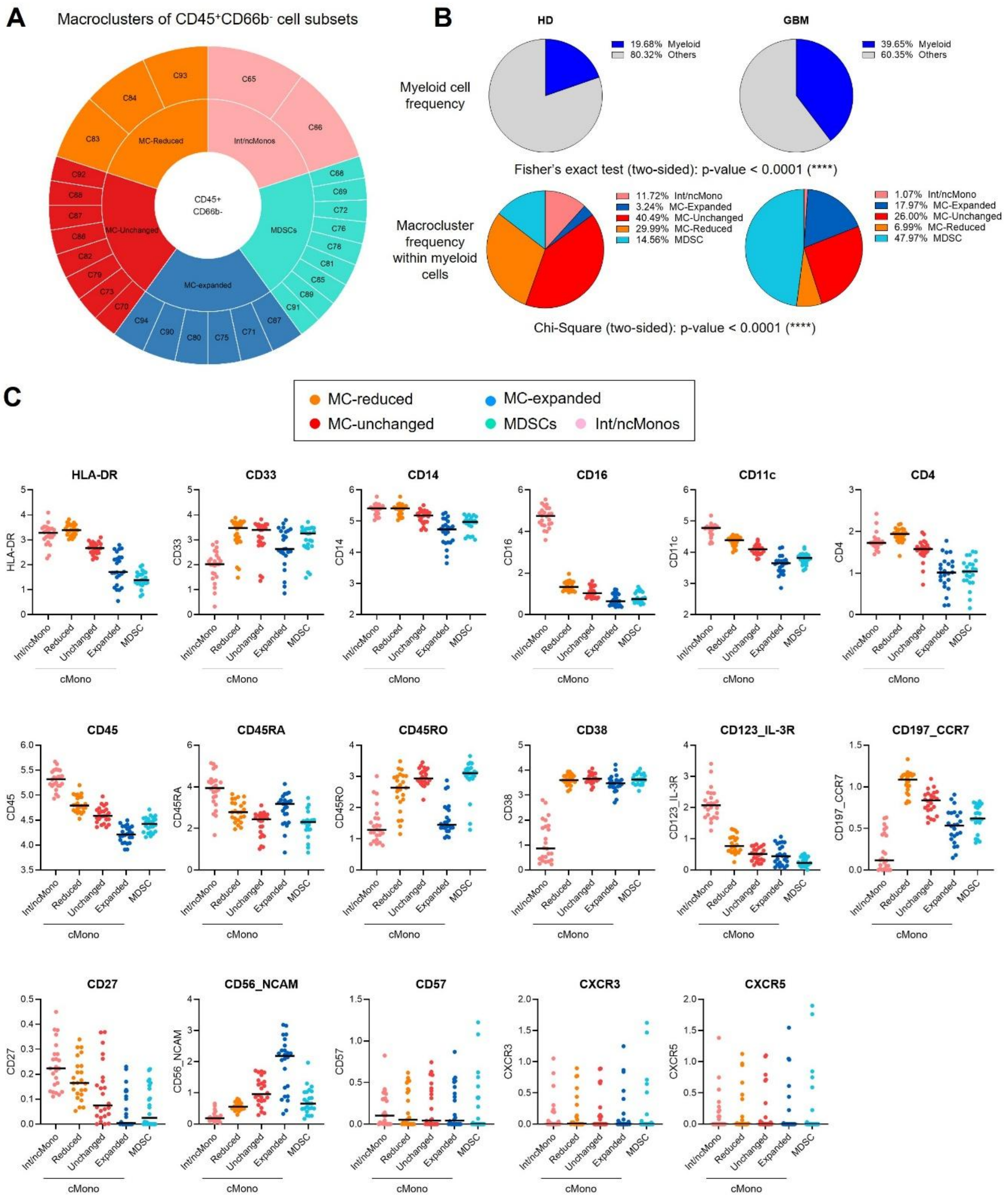


**Figure S4. Cell frequency and marker expression trends in CD45⁺CD66⁻ myeloid macroclusters.** (**A**) Macroclusters of CD45⁺CD66⁻ myeloid cell subsets obtained by grouping classical monocytes (cMonos) and monocytic dendritic cells (mDCs), intermediate and non-classical monocytes (int/ncMonos) and MDSCs. Clusters of cMonos and mDCs were further categorized as expanded, reduced or unchanged in GBM patients relative to HDs, based on their modulation trends from supervised analysis. MC: macrocluster; Int/ncMonos: intermediated/non-classical monocytes. (**B**) Frequency of myeloid cells in HDs or GBM patients, expressed as proportion of total cells (top pie charts) or as proportion of overall myeloid compartment (bottom pie charts). Contingency analyses revealed significant modulation in frequency and across macroclusters in GBM patients compared with HDs (myeloid cell frequency: Fisher's exact test, p-value < 0.0001; macrocluster frequency within myeloid cells: Chi-square tests; p-values < 0.0001). (**C**) Trends of cell surface marker expression across the five myeloid macroclusters, corresponding to the analysis of **Fig. 2B**.

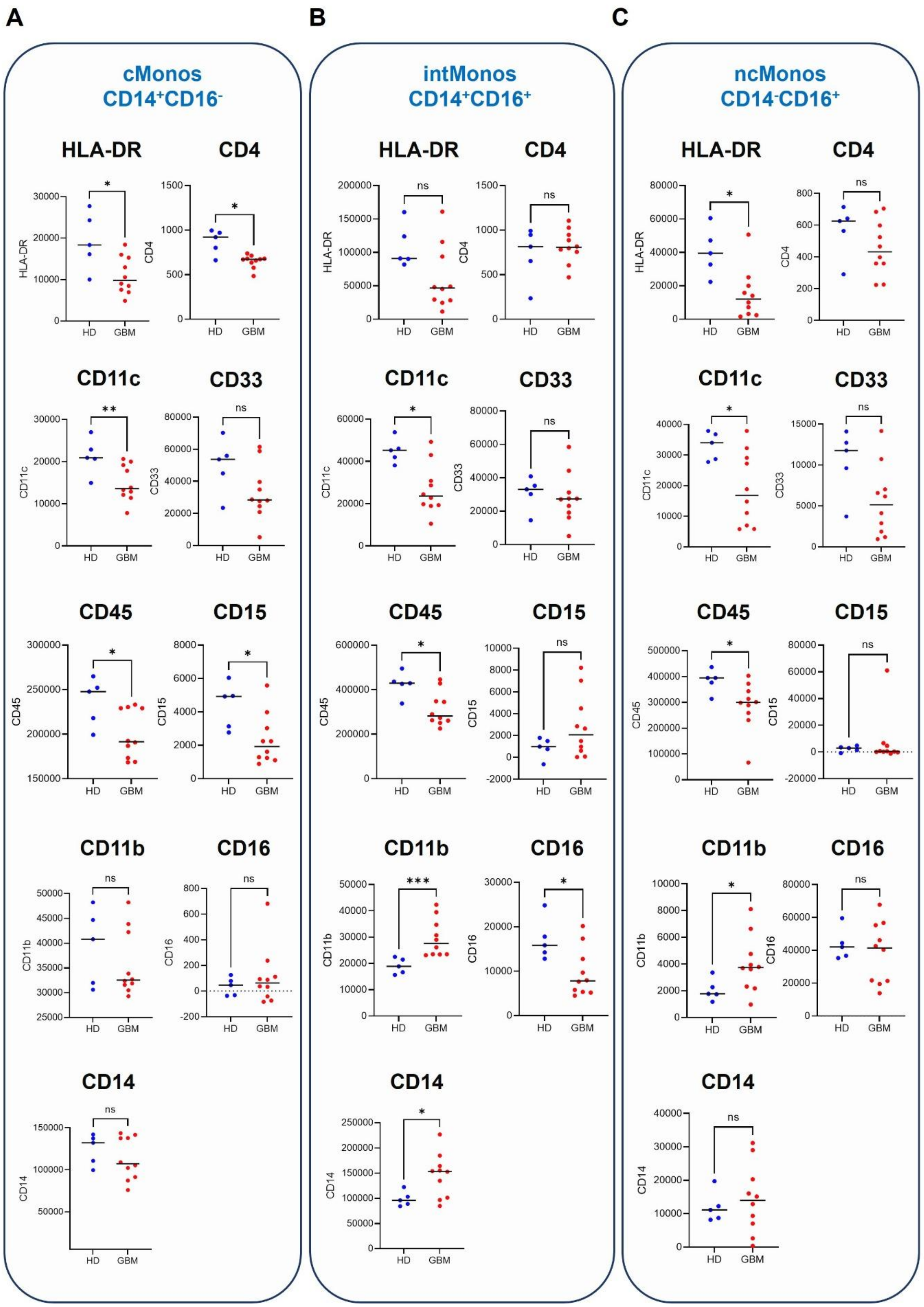


**Figure S5. Flow cytometry quantification of selected markers in classical, intermediate and non-classical monocytes.** Expression of selected cell surface markers (arbitrary units; median values) in (**A**) classical monocytes (cMonos), (**B**)

intermediate monocytes (intMonos) and (**C**) non-classical monocytes (ncMonos) from HDs (n=5) and GBM patients (n=10). Non-parametric Mann-Whitney U test (*: $p < 0.05$; **: $p < 0.01$; ***: $< 0.001$; ns: not significant).

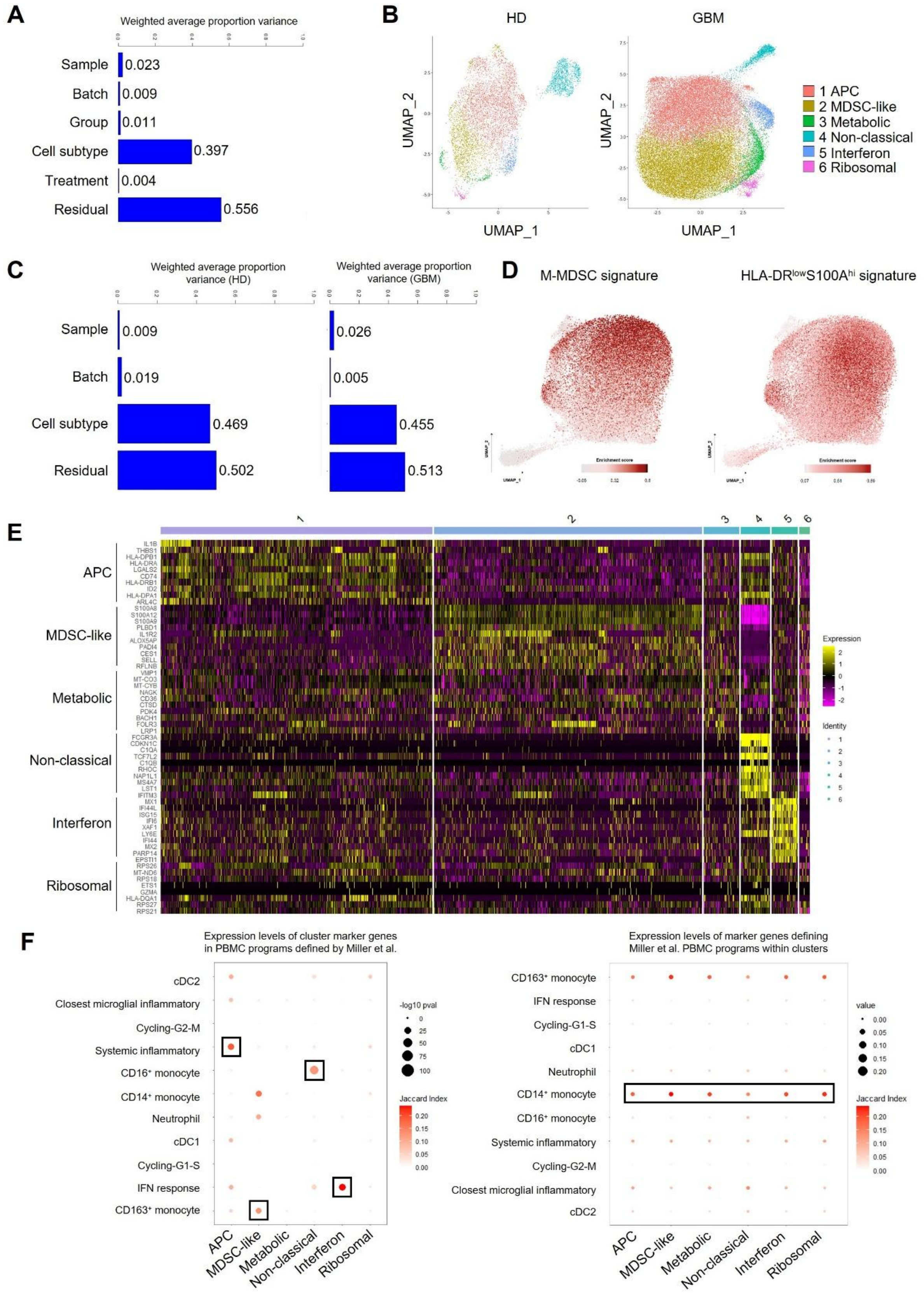

**Figure S6. Characterization of monocytic transcriptional subsets.** (**A**) Bar chart showing the contribution of the co-variates to the weighted average proportion variance underlying monocytic transcriptomic heterogeneity in HDs and GBM patients. (**B**) UMAPs showing the six monocytic clusters in HDs (n=5; left) or GBM patients (n=10; right). (**C**) Bar chart showing the contribution of the co-variates to the weighted average proportion variance underlying monocytic transcriptomic heterogeneity in HDs (left) or GBM patients (right). (**D**) UMAPs showing the identity of the recognised MDSC-like cluster using literature-based MDSCs and HLA-DR$^{low}$S100A$^{high}$ transcriptional signatures. (**E**) Heatmap depicting top up-regulated genes (adjusted p-value < 0.05, Avg_Log$_2$FC ≥ 0.25; **Table S7**) characterizing each monocytic subset. (**F**) Left: similarity between monocyte cluster markers and top 100 genes underlying PBMC programs defined by Miller and colleagues. Right: expression levels of marker genes defining Miller et al. PBMC programs within clusters.

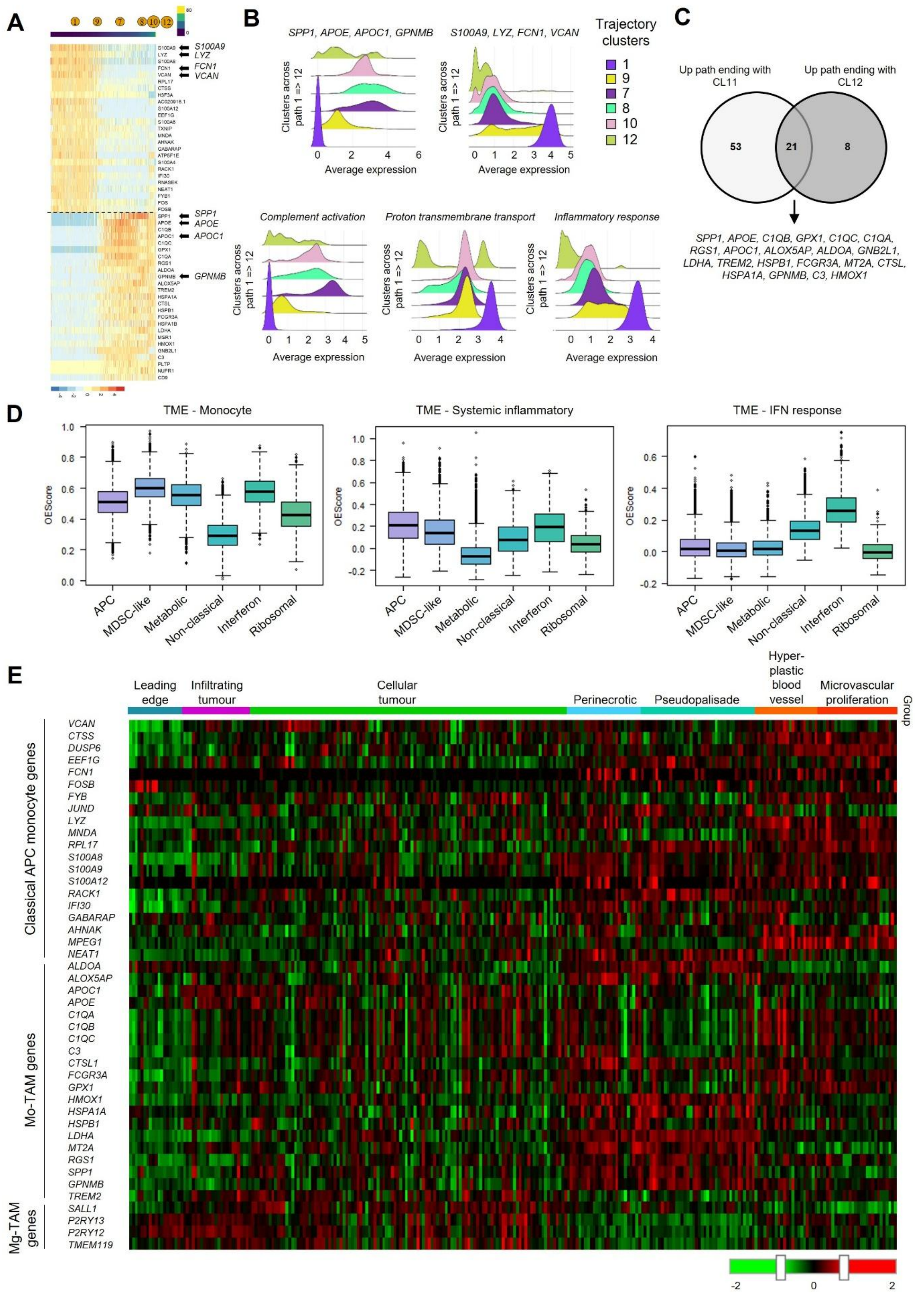


**Figure S7. Monocyte subsets undergo specific differentiation trajectories toward tumour-associated macrophages and display specific spatial localization within the tumour.** (**A**) Heatmap showing the top 20 downregulated (top) and downregulated (bottom) genes along the trajectory from cluster 1 to cluster 12 (**Table S11**). (**B**) Top: ridge plots showing the average expression level of prototypical glioma-associated macrophage markers (*SPP1, APOE, APOC1* and *GPNMB*) and monocytic-specific genes (*S100A9, LYZ, FCN1* and *VCAN*) along the trajectory from cluster 1 to 12. Bottom: ridge plots representing the average expression level of GO specific gene signatures along the trajectory from cluster 1 to 12: complement activation (GO:0006958; *C1QB, C3, C1QA, C1QC*), proton transmembrane transport (GO:1902600; *MT-ATP6, RNASEK, MT-CO1, MT-CO2, MT-CO3, MT-ND3, ATP6V0C, ATP5F1E, MT-ND1*) and inflammatory response (GO:0006954; *ANXA1, NAMPT, CYBB, TXNIP, S100A12, FOS, LYZ, S100A9, S100A8*). (**C**) Venn diagram showing shared and distinct up-regulated genes between trajectories ending with cluster 11 (CL11) and 12 (CL12). (**D**) Expression levels of TME programs defined by Miller and colleagues across monocytic subsets. (**E**) Heatmap showing expression levels of selected classical APC monocyte genes and Mo-TAM genes modulated across the monocytic differentiation into Mo-TAMs and Mg-TAM genes extracted from the Ivy Glioblastoma Atlas Project dataset across laser dissected specific tumour regions: leading edge, infiltrating tumour, cellular tumour, perinecrotic, pseudopalisade, hyperplastic blood vessel and microvascular proliferation areas. Colour bar denotes Z-scored gene expression levels.

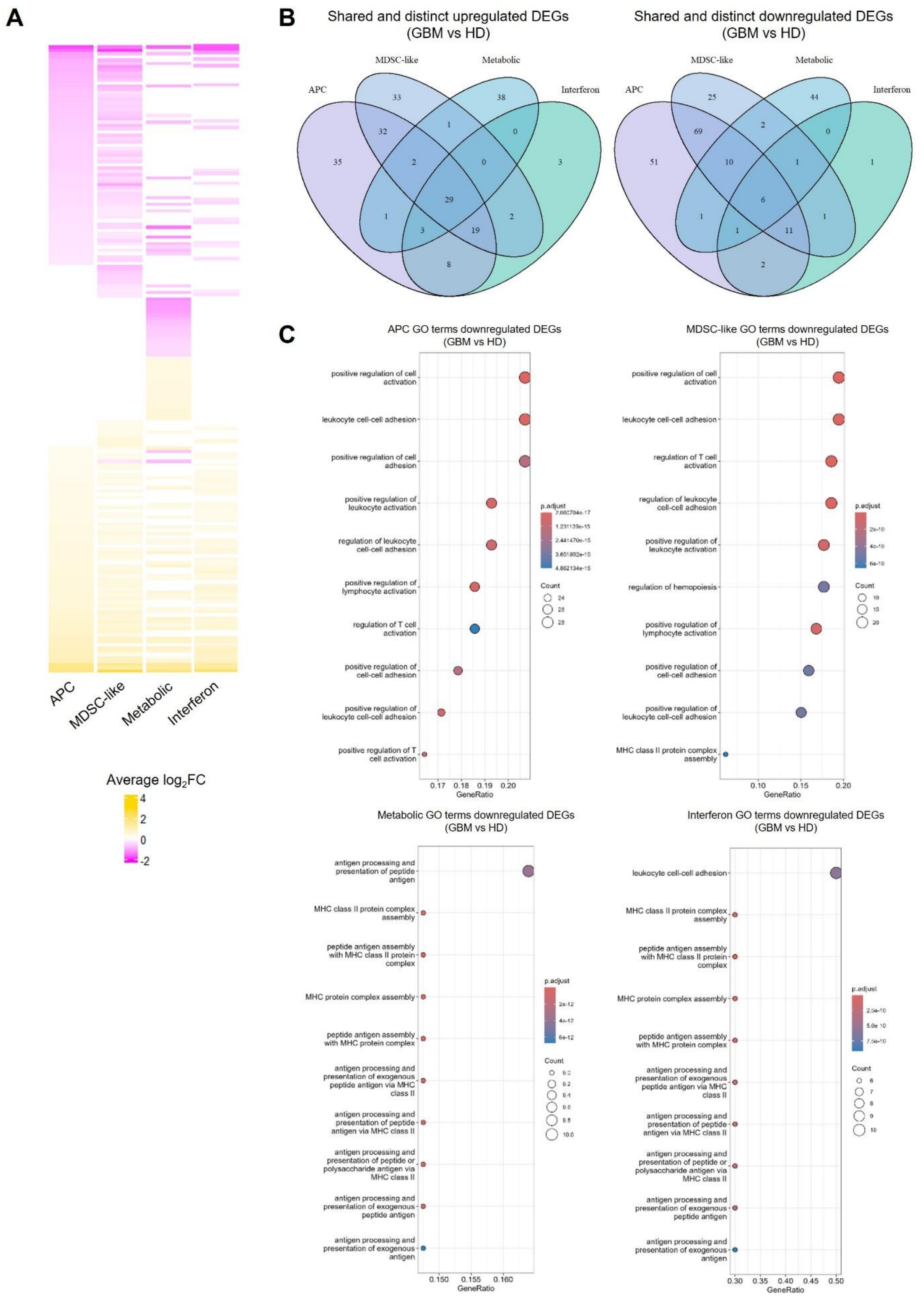


**Figure S8. Characterization of classical monocytic clusters.** (**A**) Heatmap depicting all DEGs between GBM patients and HDs (adjusted p-value < 0.05, |Avg_log$_2$FC| ≥ 0.25; **Table S12**) across APC, MDSC-like, metabolic and interferon monocytic subsets. Colour bar denotes average $log_2FC$. (**B**) Venn diagrams showing number of shared and discrete up-regulated (left) and down-regulated (right) genes across four classical monocyte subsets. (**C**) Dot plots depicting main GO biological processes (BP) associated with down-regulated genes comparing GBM patients and HDs in the identified four monocytic clusters (**Table S13**). Colour bar denotes adjusted p-value and circle diameter depicts gene count of modulated genes within the GO-BP term.

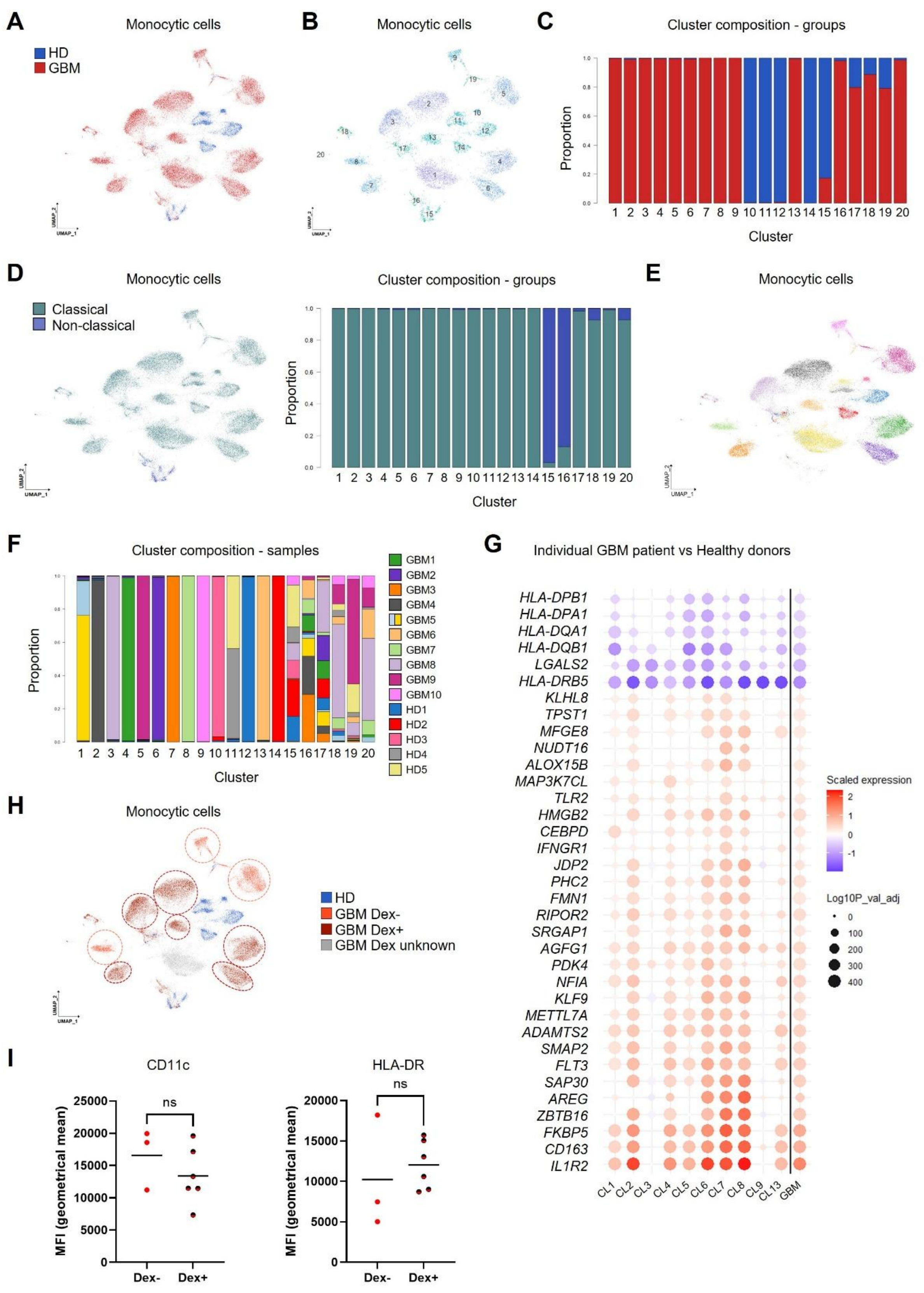


**Figure S9. GBM patient-derived classical monocytes show inter-individual differences at the transcriptomic level.** (**A**) UMAP showing re-clustered monocytes from UMAP depicted in **Fig. 3C** prior batch correction. (**B**) UMAP showing 20 welldefined monocytic clusters. (**C**) Bar chart displaying the proportion of cells derived either from GBM patients or HDs in each monocytic cluster. (**D**) Left: UMAP showing the discrimination between classical (grey) and non-classical (purple) monocytic cells. Right: Bar chart displaying the proportion of either classical or non-classical monocytes in each monocytic cluster. (**E**) UMAP depicting cell distribution derived from each individual across monocyte clusters. (**F**) Bar chart showing the proportion of cells derived from each individual across all the monocyte clusters. (**G**) Dot plot showing average fold change expression of GBM-CMS genes in each patient-specific classical monocyte cluster. Circle diameter denotes $\log_{10}$ (adjusted p-value) and colour depicts average fold change of each gene between cluster cells and HD-derived classical monocytes. (**H**) UMAP displaying monocytic cells from HDs (blue) and from GBM patients either treated (dark red), untreated (light red) with dexamethasone, or with unknown treatment status (grey). (**I**) Dot plot showing geometrical mean fluorescent intensity (MFI) analysis by multicolour flow cytometry of CD11c and HLA-DR in $CD14^+CD16^-$ classical monocytes between GBM patients untreated (GBM dex-, n=3) or treated (GBM dex+, n=6) with dexamethasone. Unpaired t-test (ns: not significant).

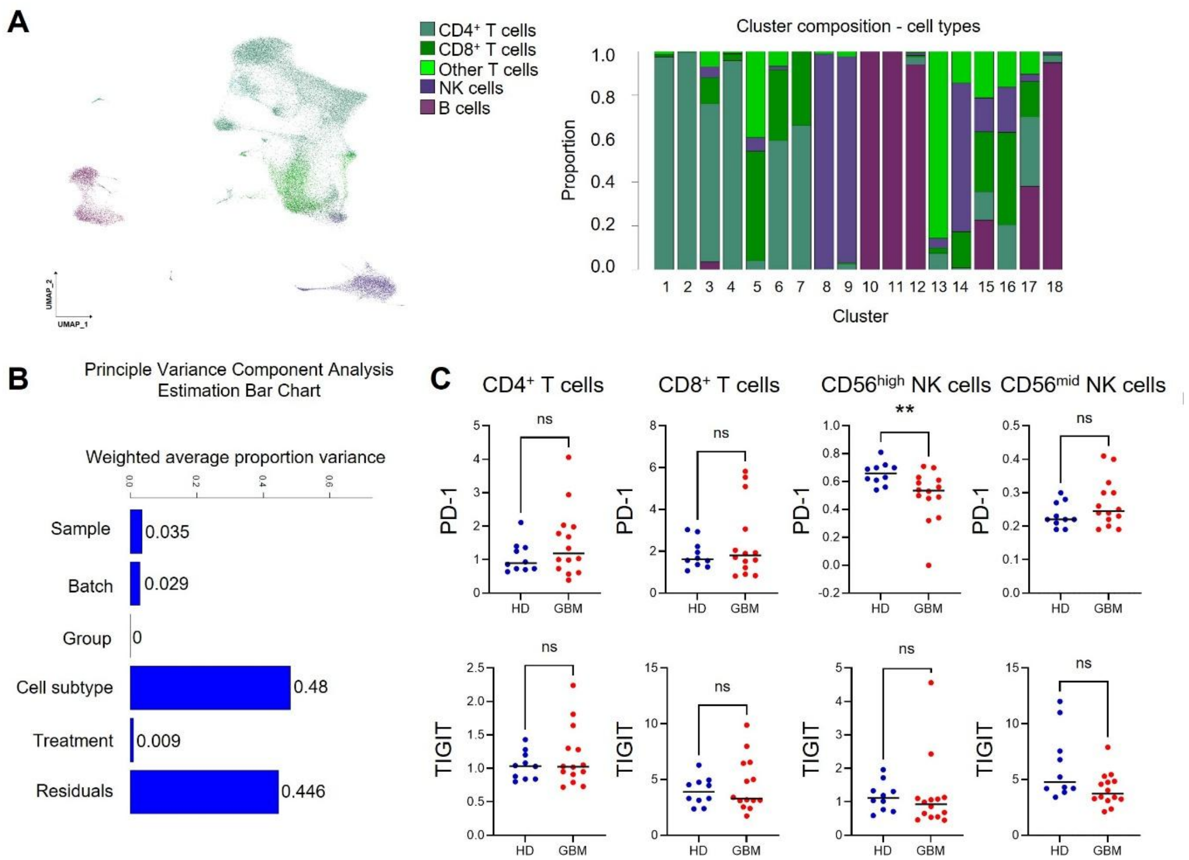


**Figure S10. Characterization of lymphocytic clusters.** (**A**) Left: UMAP showing cell type assignation from Azimuth: $CD4^+$ T cells (dark green), $CD8^+$ T cells (bright dark green), other T cells (light green), NK cells (purple) and B cells (magenta). Right: Bar chart showing the composition of cell types in each cluster. (**B**) Bar chart showing the contribution of the co-variates to the weighted average proportion variance underlying lymphocyte transcriptomic heterogeneity. (**C**) Dot plot displaying expression levels of PD-1 and TIGIT in $CD4^+$ T cells, $CD8^+$ T cells, NK $CD56^{dim}$ cells and NK $CD56^{bright}$ cells, extracted from

GBM patients and HDs determined by mass cytometry. Unpaired t-test (n=14 GBM patients, n=10 HDs; **: p-value < 0.01; ns: not significant).

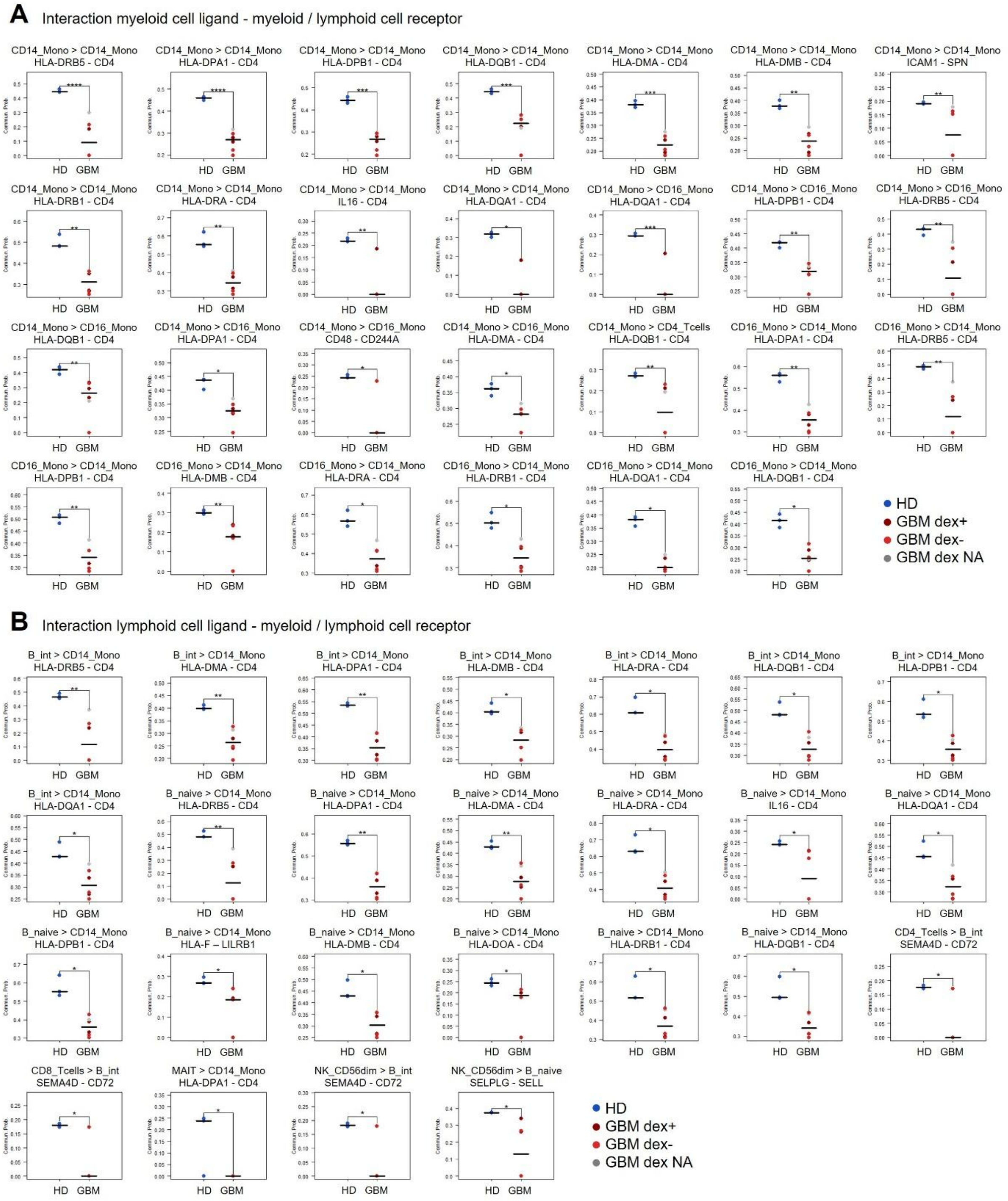


**Figure S11. Changes of ligand-receptor interactions in GBM patients are independent from dexamethasone therapy.** (**A-B**) Dot plots showing all interactions whose sender cells are (**A**) myeloid cells ($CD14^{+}$ or $CD16^{+}$ monocytes) or (**B**) lymphocytes (intermediate or naïve B cells, $CD4^{+}$ or $CD8^{+}$ T cells, MAIT and $CD56^{dim}$ NK cells) altered in PBMCs comparing

GBM patients (n=6) and HDs (n=3) (adjusted p-value < 0.05). GBM patients discriminated according to dexamethasone therapy: treated (GBM dex+, dark red, n=2), untreated (GBM dex-, light red, n=3), unknown (GBM dex NA, grey, n=1).

#### LEGENDS FOR SUPPLEMENTARY DATA FILES

**Table S1. Clinical data of glioblastoma (GBM) patients and healthy donors (HDs) included in this study**

**Table S2. Cell subset classification based on surface marker expression (median values, mass intensity units)**

**Table S3. Unsupervised (Mann-Whitney U-test; Qlucore) and supervised (cell lineage assignment) analysis on 100-metacluster resolution**

**Table S4. Comparative analysis of immune cell frequencies by unsupervised and supervised analyses (CyTOF data; Mann-Whitney U-test; Prism) and flow cytometry data (Mann-Whitney U-test; Prism)**

**Table S5. Sequential gating strategy of the CyTOF data (FlowJo analysis)**

**Table S6. Supervised cell lineage enrichment from manual sequential gating of CyTOF data (Mann-Whitney U-test; Qlucore)**

**Table S7. Genes characterising each individual monocyte cluster**

**Table S8. List of GO biological processes enriched in each monocyte cluster**

**Table S9. List of transcription factors characterising each monocyte cluster**

**Table S10. DEGs across trajectories (ending with CL4, CL5 or CL6)**

**Table S11. DEGs across trajectory (ending with CL11 or CL12)**

**Table S12. DEGs comparing GBM patients and HDs across monocytic clusters (adjusted p-value < 0.05, |Avg_log2FC| ≥ 0.25)**

**Table S13. List of GO biological processes corresponding to up- or down-regulated DEGs across monocytic clusters comparing GBM patients with HDs (adjusted p-value < 0.05)**

**Table S14. DEGs comparing GBM patients and HDs across lymphocytic clusters (adjusted p-value < 0.05, |Avg_log2FC| ≥ 0.25)**

**Table S15. Signalling pattern comparison between GBM patients and HDs (adjusted p-value < 0.05)**

**Table S16. Ligand-receptor comparison between GBM patients and HDs (adjusted p-value < 0.05)**

**Table S17. Materials and methods**
